# Age-related decline of PKA-RIIβ level in SNc dopaminergic neurons underlies PD pathogenesis

**DOI:** 10.1101/2024.09.13.612981

**Authors:** Yun Zhao, Bingwei Wang, Miao Zhao, Jiarui Liu, Yujia Hou, Shizhang Wei, Changxian Xiong, Daotong Li, Ruimao Zheng

**Affiliations:** Department of Anatomy, Histology and Embryology, School of Basic Medical Sciences, Peking University, Beijing, China; Beijing Life Science Academy, Beijing, China; Department of Biomedical Informatics, Center for Noncoding RNA Medicine, School of Basic Medical Sciences, Peking University, China; Neuroscience Research Institute, Peking University, Beijing, China; Key Laboratory for Neuroscience of Ministry of Education, Peking University, Beijing, China; Key Laboratory for Neuroscience of National Health Commission, Peking University, Beijing, China

## Abstract

The cyclic-AMP dependent protein kinase A (Protein kinase A, PKA) regulates dopaminergic function in the substantia nigra pars compacta (SNc). However, whether PKA is involved in the pathogenesis of Parkinson’s disease (PD) is unknown. Here, by collecting and analyzing the current worldwide SNc transcriptomic datasets of PD patients, we found a decline of PKA-RIIβ subunit level in the SNc of PD patients. The decreased PKA-RIIβ subunit level was positively correlated with decreased dopamine synthesis and increased oxidative stress in the SNc of PD patients. PKA-RIIβ subunit is expressed in the striatum and the SNc. PKA-RIIβ gene knockout mice (RIIβ-KO) showed a age-related parkinsonism at 12 months of age. Using Cre-LoxP system, we observed that RIIβ re-expression in the SNc dopaminergic neurons rescued parkinsonism of RΙΙβ-KO mice. RIIβ re-expression in striatal neurons did not affect parkinsonism of RΙΙβ-KO mice. The spontaneous parkinsonism could be developed in 12-month-old SNc dopaminergic neuron-specific RIIβ-deficient mice. Single-nucleus RNA sequencing revealed decreased PKA activity, reduced dopamine synthesis and raised oxidative stress in the SNc dopaminergic neurons of RIIβ-KO mice. Adeno-associated virus (AAV)-mediated gene therapy targeting PKA-RΙΙβ in the SNc dopaminergic neurons rescued parkinsonism in PD mouse model. Taken together, these findings indicate that PKA-RIIβ may be a key factor of human genetic etiologies of PD. The therapy targeting PKA-RIIβ in the SNc dopaminergic neurons may be promising for PD treatment.

## Introduction

Parkinson’s disease (PD) is the second most common neurodegenerative disorder, affecting approximately 1 % of the individuals over age 60 years and 4% of the population older than age 85^1–5^. Clinical features of PD include rest tremor, rigidity, and bradykinesia^6–8^. Pathology of PD is mainly characterized by decreased dopamine synthesis, increased oxidative stress, and loss of dopaminergic neurons in the substantia nigra pars compacta (SNc) of midbrain^9–13^. The cause of PD remains unknown.

Since 1971, pharmacological studies have revealed that the cyclic-AMP (cAMP) dependent protein kinase A (Protein kinase A, PKA) is essential for the function of SNc dopaminergic neurons^14–21^. PKA holoenzyme is a heterotetramer consisting of two regulatory (R) and two catalytic (C) subunits^22–24^. In humans, four R subunit genes (encoding RIα, RIβ, RIIα, and RIIβ) and two C subunit genes (encoding Cα and Cβ) have been identified^25–27^. PKA holoenzyme is inactive^28^. The binding of cAMP to the R subunits releases active C subunits to phosphorylate downstream substrates^29^. The R subunits also act as intrinsic inhibitors of the C subunits, and protect the C subunits from degradation^30,31^. PKA is involved in the dopamine biosynthesis and oxidative stress in the dopaminergic neurons^32–35^. In the SNc dopaminergic neurons, decreased activity of tyrosine hydroxylase (TH), the rate-limiting enzyme of dopamine synthesis, leads to dopamine deficiency and dysfunction in the striatum (STR)^36–38^. PKA regulates TH activity by phosphorylating it at serine 40^39–45^. Altered cytoplasmic phosphorylated cAMP-response element binding protein (CREB), a pivotal PKA-phosphorylated transcription factor, was observed in the SNc dopaminergic neurons of PD patients^46,47^. PKA activators effectively rescue PINK1 knockdown-induced mitochondrial dysfunction and cell death in human dopaminergic cells^17,47^. Notably, the PKA-RIIβ subunit is highly expressed in human or murine SNc and striatum^48,49^. The RIIβ gene knockout mice exhibit altered movement behavior^48^. Nevertheless, whether RIIβ-PKA in the SNc dopaminergic neurons is linked to PD pathogenesis in unknown.

In this study, by using the current worldwide human SNc transcriptomic datasets, we detected a remarkably lowered RIIβ subunit level in the SNc of PD patients. By applying multiple genetically engineered mouse models and single-nucleus RNA sequencing (snRNA-seq) technology, we found that an age-related decline of PKA-RIIβ level in the SNc dopaminergic neurons may be a culprit in PD. RIIβ subunit-targeted gene manipulation effectively relieved PD-like symptoms in mice. In summary, we suggest that RIIβ-PKA in the SNc dopaminergic neurons may act as a key contributor to PD pathogenesis. RIIβ-PKA may be a promising target in gene therapy for PD.

## Methods

### Mice and animal Care

C57BL/6 mice were purchased from Charles River Laboratories Beijing Branch (Beijing Vital River Laboratory Animal Technology Co., Ltd.) and the Department of Laboratory Animal Science of Peking University Health Science Center. All experiments were performed on C57BL/6 male mice (the age of the mice was described in the study). Age-matched mice were used for each experiment. All procedures were reviewed and approved by the Institutional Care and Use Committee of the Peking University Health Science Center (approval number: LA2019340). Mice were housed at 22 ± 1 °C with a 12-hours light/dark cycle (light time, 20:00 to 8:00) with unrestricted access to food and water. RIIβ knockout (RIIβ^−/−^) mice and RIIβ^lox/lox^ mice were a kind gift from Dr. G. Stanley McKnight of University of Washington^49,50^. D1R-Cre mice were a kind gift from Dr. Yi Rao of Peking University^51^. A2A-Cre mice and DAT-Cre mice were a kind gift from Dr. Peng Cao of National Institute of Biological Sciences^52,53^. Genotyping of mice carrying mutated RIIβ, lox RIIβ allele, D1R-Cre, A2A-Cre and DAT-Cre were performed as previously described. To minimize the occurrence of germ-line recombination, heterozygous DIR-RIIβ knocked (D1R-Cre/RIIβ^+/−^) mice were crossed with RIIβ^lox/lox^mice to generate heterozygous RIIβ DIR-Cre mice (D1R-Cre/RIIβ^lox/−^); heterozygous A2A-RIIβ knocked (A2A-Cre/RIIβ^+/−^) mice were crossed with RIIβ^lox/lox^ mice to generate heterozygous RIIβ A2A-Cre mice (A2A-Cre/RIIβ^lox/−^); heterozygous DAT-RIIβ knocked (DAT-Cre/RIIβ^+/−^) mice were crossed with RIIβ^lox/lox^mice to generate heterozygous RIIβ DAT-Cre mice (DAT-Cre/RIIβ^lox/−^). RIIβ^flox/flox^ mice were generated by using CRISPR/Cas9 technology at Biocytogen. The RIIβ^flox/flox^ mice were crossed with D1R-Cre mice to obtain D1R-Cre/RIIβ^flox/flox^ mice; the RIIβ^flox/flox^ mice were crossed with A2A-Cre mice to obtain A2A-Cre/RIIβ^flox/flox^ mice; the RIIβ^flox/flox^ mice were crossed with DAT-Cre mice to obtain DAT-Cre/RIIβ^flox/flox^ mice. During all procedures of experiments, the number of animals and their suffering by treatments were minimized.

### MPTP administration

Mice were injected intraperitoneally l-methyl-4-phenyl-l,2,3,6-tetrahydropypridine (MPTP; 30 μmg/kg; Sigma-Aldrich) for 5 days, as previously described^54,55^, and the controls were received an equivalent volume of 0.9 % saline.

### Immunohistochemistry and immunofluorescence

Primary antibodies and working dilutions are detailed in Supplementary Table 1. Immunohistochemistry and immunofluorescence were performed on 30-μm thick serial brain sections. Either mouse was perfused transcardially and brains fixed in 4 % paraformaldehyde (PFA) for 2 days. Brains were transfer to 20–30 % gradient sucrose for cryoprotection, followed by freezed in OCT compound (Sakura FineTech, Tokyo). For histological studies, brain slices were blocked with 10 % goat serum in PBS with 0.2 % Triton X-100 and incubated with TH antibody overnight at 4 □. After washing with PBS three times, brain sections were incubated with biotin secondary antibody, followed by avidin-biotin complex (Zsbio, SP-9001) and visualized with 3,3’-diaminobenzidine (DAB) peroxidase substrate (Zsbio, ZLI-9018). For immunofluorescence, brain pieces were blocked with 10 % goat serum in PBS with 0.2 % Triton X-100 and incubated with TH or RIIβ antibodies overnight at 4 □. After washing with PBS three times, brain sections were incubated with secondary antibodies (Alexa-Fluor 488- or 594-conjugated secondary antibodies, 1:400, YEASEN) for 2 hours at room temperature. Images were captured on a Leica microscope (DMI 4000B, Wetzlar, Germany) or Olympus VS120 Virtual Microscopy Slide Scanning System The selected area in the signal intensity range of the threshold, the number of TH- or RIIβ-positive neurons in the SNc, and the density of TH positive fibers in the striatum were measured using Image J software.

### Protein extraction and western analysis

Proteins were extracted from SNc or STR using ice-cold RIPA lysis buffer [0.5 % NP-40, 0.1 % sodium deoxycholate, 150 mM NaCl, 50 mM Tris-HCl (pH 7.4), phosphatase inhibitors (B15002, Bimake), and protease inhibitor cocktail (B14002, Bimake)]. Lysates were centrifuged at 12,000 g for 30 minutes at 4 °C. Supernatants from lysates were used as protein extracts, protein concentration was calculated by BCA assay (Aidlab; PP01). Protein extracts was added protein loading buffer [62.5 μmM Tris•Cl (pH 6.8), 2 % (wt/vol) SDS, 5 % glycerol, 0.05 % (wt/vol) bromophenol blue] and denatured by boiling at 95 °C for 5 minutes. Equal amounts of proteins were separated by 10 % sodium dodecyl sulfate poly acrylamide (SDS-PAGE), transferred to nitrocellulose membrane (Pall Corporation; T60327). The membranes were blocked in 5 % skim milk (Tris-buffered saline and Tween 20, TBST) for 2 hours at room temperature, followed by incubation with the primary antibody in 5 % BSA-TSBT overnight at 4 °C. After overnight incubation, the membrane was washed three times in TBST for 15 minutes, and then incubation with HRP-conjugated secondary antibody in TBST with 5 % skim milk for 2 hours at room temperature. Following three cycles of 15 minutes washes with TBST, the membranes were developed using an Enhanced Chemiluminescence assay (BIO-Rad). Intensities of the protein bands in western blot images were analysis by the ImageJ software (NIH). All raw blot images are available in sFigure 8, sFigure 9 and sFigure 10.

### Behavioral tests

To evaluate the motor performance of mice, beam traversal, pole test, rotarod test, hindlimb scoring and gait test were used. The experimenter was blind to the genotype of mice. Before the first trial of the day, mice were acclimated to the test room for 60 minutes and 1 day prior to testing. Before the actual test, mice were trained for three consecutive days. Animals were given 3 trials, each separated by 10 minutes. Apparatus was cleaned with 75 % ethanol on each trial.

Beam traversal test. Beam test was used to detect subtle deficits in motor skills and balance of mice. A 100 cm wooden beam consists of four segments of 0.25□m in length. Each segment was of thinner widths 3.5□cm, 2.5□cm, 1.5□cm, and 0.5□cm, with 1□cm overhangs placed 1□cm below the surface of the beam. In the test, mice were placed on the widest segment as a loading platform, the narrowest segment placed into a dark goal box. Mice were made to traverse the beam in the same manner (cut-off time 30 seconds maximum). The test time from the start to the 90 cm point was recorded. Timing began when the animals placed their forelimbs onto the 2.5□cm segment and ended when one forelimb reached the 90 cm point.

Pole test. The pole test is conducted on a wooden rod (diameter 8 mm; height 80 cm) which was wrapped with bandage gauze. The rod was fixed in the middle of an empty cage. Mice were placed the top of a wooden pole and facing downward. The test time until it descended to the base of the pole was recorded with a maximum duration of 30 seconds. When the mouse was not able to turn downward and instead dropped from the pole, the test time was taken as the slowest mouse to pass the pole. The pole test is used to assess rigidity.

Rotarod test. Rotarod test evaluates motor coordination and motor learning of mice. In the test, mice were placed on an the accelerating rotarod cylinder. After pre-training at 4 rpm for one minute, the speed was gradually increased from 4 to 40 rpm within 5 minutes, and kept at 40 rpm for additionally 2 minutes. A trial ended if mice fell off the rungs or gripped the device and spun around for two consecutive revolutions without attempting to walk on the rungs. Time before falling was automatically recorded with a maximum duration of 5 minutes. Data are presented as the percentage of the 3rd trials on the rotarod compared to the control.

Hindlimb scoring. Mice were gently lifted upward by the mid-section of the tail and observed over 5–10 seconds. Mice were assigned a score of 0, 1, 2, 3 based on the extent to which the hindlimbs clasped inward. The mice that freely moved and extended their limbs outward were scored as 0. A score of 1 was recorded if the mice kept one hindlimb inward while restrained or showed partial inward clasping with both legs. A score of 2 was assigned when both legs were clasped inward for most of the observation period, but still exhibited some flexibility. If mice exhibited full hindlimb paralysis with immediate inward clasping and no flexibility, a score of 3 was assigned.

Gait test. The testing apparatus is constructed from a 3 mm thick gray acrylic board and includes a runway with non-slippery white paper (10 cm wide, 60 cm long, 12 cm tall) and a dark goal box (16 cm wide, 10 cm long, 12 cm tall). During first training day, mice were familiarized with the equipment for 2 minutes before having their front and back paws colored red and black using safe food dyes. Mice were then trained to run to the goal box. In the test, mice were required to run the runway within a maximum time of 60 seconds. The analysis of footprint patterns focused on three parameters (stride length, stride width, and overlap), with prints near the beginning and end disregarded due to the impact of acceleration or deceleration. Stride length was measured as the average distance between each forepaw and hindpaw footprint. Stride width was measured as the average distance between the right and left footprint of each forepaw and hindpaw. At least four values were measured in each trial for each parameter.

### Stereotaxic Injection of AAV

Mice were anesthetized with isoflurane and placed in a stereotaxic apparatus. The mice were placed in a stereotaxic holder to stabilize brains and the skull was exposed via a small scalp incision. By using a 0.2□mm-gauge stainless steel injector connected to a 5 μl Hamilton syringe, AAV2/9-EGFP, AAV2/9-hSyn-RIIβ-S112A and AAV2/9-Flex-RIIβ-S112A (2 × 10^12^ viral particles) were bilaterally injected into the SNc region (coordinates, bregma: anterior-posterior, −3.2 mm; dorsal-ventral, ±1.2 mm; lateral, −4.6 mm, according to the atlas of Paxinos and Franklin). Only animals with correct injection placements, verified by analyzing immunofluorescence staining of consecutive coronal brain sections, were included in the statistical analysis.

### Bulk RNA-seq and analysis

SNc tissues were obtained from three male RIIβ-KO mice and three male WT mice (12 months old). The total RNA from the SNc was extracted using TRIzol reagent (TransGen Biotech). A total amount of 3 µg RNA per sample was used as input material for the RNA sample preparations. After that, the RNAs were subjected to 50 bp single-end sequencing with a BGISEQ-500 sequencer. At least 20 million clean reads of sequencing depth were obtained for each sample. Differential expression analysis was performed using the limma R package (version 3.46.0). Limma package utilizes a linear model to evaluate the differentially expressed genes (DEGs). The resulting p values were adjusted using Benjamini and Hochberg’s approach for controlling the false discovery rate. Genes with an adjusted *p* < 0.05 found by limma package were assigned as differentially expressed. DEGs were defined as genes with FDR less than 0.01 and log2 fold change larger than 1 (upregulation) or smaller than −1 (downregulation). Gene Ontology (GO) and Kyoto Encyclopedia of Genes and Genomes (KEGG) analysis for these DEGs were based on the Metascape database (http://metascape.org/gp/index.html#/main/step1). The expression heatmap of all DEGs was plotted using pheatmap R package (version 1.0.12).

### snRNA-seq and analysis

Brain slice preparation for microdissection and single-nucleus dissociation. Single-nucleus dissociations for the snRNA-seq experiments were performed on SNc tissue from mouse brain slices. The snRNA-seq was as independent experiment. For snRNA-seq experiment, SNc tissues were obtained from five male RIIβ-KO mice (pooled) and five male WT mice (pooled) (12 months old). All mice were killed by rapid decapitation following isoflurane anesthesia, within the same time (morning, 8:00–11:00). Three slices were obtained from each animal that approximately corresponded to mouse brain atlas images representing the distance from bregma −2.92, −3.16 and −3.64 mm, as indicated in Figure 5A. These slices were homogenized in homogenization buffer (250□mM sucrose, 5□mM MgCl2, 25□mM KCl, 10□mM Tris buffer, 1□μM DTT, protease inhibitor, 0.4□U/μL RNaseIn, 0.2□U/μL Superasin, 0.1% Triton X-100, 1□μM propidium iodide and 10□ng/mL Hoechst 33342).

Nucleus capture and sequencing. After being filtered through 40 μm nylon mesh filters to remove any cell aggregates, the samples were centrifuged at 3000 g for 8 minuts at 4 °C, and then resuspended in PBS supplemented with 0.3 % BSA, 0.4□U/μL RNaseIn and 0.2□U/μL Superasin. Nuclei that were positive for both Hoechst 33342 and PI were isolated through fluorescence-activated cell sorting (BD Influx) and quantified using a dual-fluorescence cell counter (Luna-FL™, Logos Biosystems). Following the standard protocol, approximately 12,000 nuclei for RIIβ-KO group and 8,000 nuclei for control group were isolated for each sample using the 10× Genomics Chromium Single Cell Kit v3.1 and then sequenced in the Illumina NovaSeq 6000 PE 150. For FASTQ generation and alignments, Illumina basecall files (*.bcl) were converted to FASTQs using Cell Ranger v6.1.2 (10× Genomics), which uses bcl2fastq v.2.17.1.14. FASTQ files were then aligned to mm10 genome and transcriptome using the Cell Ranger v6.1.2 software, which generates a “gene versus cell” count matrix.

**s**nRNA seq analysis. For filtering and unsupervised clustering, the gene expression matrix from Cell Ranger was applied for downstream analysis (using R 4.3.0). Filtering, integration, clustering and identification of cell types were processed under the corresponding pipeline of Seurat v4.3 package. Of the initial 18148 nucleus (7118 for WT group and 11030 RIIβ-KO group), cells with less than 500 UMIs or > 20 % of mitochondrial reads were discarded. The nucleus gene expression that remained was normalized based on the total number of transcripts identified in each nucleus and then multiplied by the median transcript count. Count matrix of each sample was normalized using the ‘LogNormalize’ function. The top 2000 genes with the most variance was identified based on their mean expression in the population and dispersion (variance/mean expression). Count matrix was scaled using the ‘ScaleData’ function. Principal component analysis (PCA) was performed with the ‘RunPCA’ function, and principal component (PC) significance was calculated using the ‘JackStraw’ function. Then dataset was clustered using the ‘FindNeighbors’ and ‘FindClusters’ function. In this case, we chose the top 25 significant PCs for downstream cluster identification and visualization. By using UMAP’s dimensionality reduction algorithm with default parameters, cell dimensionality was reduced. The marker genes of each cell type were calculated using the ‘FindAllMarkers’ function with the cutoff of |LogFC| > 0.25 and adjusted *p*< 0.05 using t test. To differentiate neuronal and non-neuronal clusters, the entire count matrix was filtered and clustered as described above. To classify a cluster as a neuron or a non-neuron cluster, the median expression of the known neuron markers syt1, snap25, syp, and eno2 were aggregated for each cluster. Every cell was then classified as a neuron or non-neuron based on their cluster membership. Subsequent clustering of non-neuronal and neuronal populations was based on this classification. For the classification of dopaminergic, GABAergic and glutamatergic clusters, clusters classified as neurons were combined and re-clustered as described above, which yielded 14 clusters. Genes differentially expressed in a given cluster were computed using limma package. GO and KEGG analysis for these DEGs were based on the Metascape database. The expression heatmap of all DEGs was plotted using pheatmap R package (version 1.0.12).

### Meta-analysis and differential expression analysis of substantia nigra transcriptome datasets of PD patients

We used ‘Parkinson’s disease’ as keywords to search for genome-wide expression studies in NCBI-GEO (http://www.ncbi.nlm.nih.gov/geo/). The inclusion criteria included as follows: (1) Only studies with sample size ≥5 in each group were included. (2) The studies that were designed for exploring gene expression in the SN for PD patients and non-PD individuals were the first choice for inclusion. A total of eight datasets were identified (GSE7621, GSE8397, GSE20333, GSE20141, GSE20292, GSE49036, GSE24378, GSE20163, GSE20164), details of the datasets are provided in Supplementary Table 2. Log2-transformation of each dataset was applied as needed. The raw CEL files were processed using the Robust Multichip Average (RMA) method for background correction and normalization, which is implemented in the affy (version 1.68.0) and gcrma (version 2.62.0) R packages. After removing duplicated gene probes and unspecific probes, all probes were mapping to single Entrez Gene IDs according to the corresponding probe annotation files. Batch effects were supervised by principal component analysis (PCA) method and removed using the ComBat function of the sva R package (version 3.38.0). Selected covariates were further adjusted for by fitting the expression data with a robust linear model and taking intercept + residuals as the adjusted expression values. Within each dataset, the expression values were further standardized to μ = 0 and σ = 1, and effect sizes (Hedges’ g) of disease status were then computed for each gene. We condensed the expression profile to the gene level, keeping the probeset with the largest effect size (i.e., the smallest dataset-specific p and thus the least likely by chance) to represent the gene. We used the R/GeneMeta package to combine effect sizes across datasets with a random effect model, compute meta-Z-score statistics, and estimate false discovery rate (FDR) by 1000 permutations.

Differential expression analysis was performed using the limma R package (version 3.46.0). Genes with an adjusted *p* < 0.05 found by limma package were assigned as differentially expressed. DEGs were defined as genes with FDR less than 0.01 and log2 fold change larger than 1 (upregulation) or smaller than −1 (downregulation). GO and KEGG analysis were based on the Metascape database. The expression heatmap of all DEGs was plotted using pheatmap R package (version 1.0.12).

For meta-analysis of RIIβ level in MPTP-treated mice, a total of six substantia nigra transcriptomic datasets of MPTP-treated mice were identified (GSE7707, GSE54795, GSE214542, GSE232039, GSE232039, one dataset from unpublished data^54^). Dataset processing and meta-analysis was described above.

### Gene co-expression network analysis

The global Multiscale Embedded Gene co-Expression Network Analysis (MEGENA) becomes a novel approach to construct large-scale co-expression planar filter networks and identify key regulator^56^. Batch effects were removed using Z-score transformation across all the samples within each dataset. The R software package MEGENA (v.1.3.7) was applied for MEGENA. The steps as follows: (1) Default parameters and a minimum module size of 50 was used to calculate a planar filtered network (PFN). (2) From the initial PFN of the connected components, then conducting multi-scale clustering on each parent cluster to generate more sub-modules, and finally analyzing the hierarchical clustering results. (3) Multiscale Hub Analysis (MHA) to identify highly connected hub genes of each module at each scale. (4) By utilizing Cluster-Trait Association Analysis (CTA), the correlation between module and clinical symptoms was calculated. (5) GO and KEGG analysis for modules were based on the Metascape database, *p* < 0.05 were assigned as the cut-off criterion for identifying enrichment

### Statistical analysis

Data are expressed as mean ± standard error of means (SEM). Representative morphological images were taken from at least three biologically independent experiments with similar results. Statistical significance was determined using Student’s t-test, one-way or two-way ANOVA, and then either Tukey’s or Bonferroni’s multiple comparison test to compare all treatment groups. p values were indicated with * *p* < 0.05, ** *p* < 0.01 or *** *p* < 0.001 on graphs. Sample sizes (n), statistical tests and p values are indicated in each figure legend.

## Results

### Human transcriptomic analysis shows that PKA-RIIβ may be linked to PD pathogenesis

To explore whether PKA is involved in PD pathogenesis, the current worldwide human SNc transcriptome sequencing datasets were collected and analyzed. The inclusion or exclusion criteria for datasets were shown in Figure.1A. 9 datasets, a total of 89 SN samples of PD patients and 79 SN samples of non-PD individuals were included. A total of 119 datasets dropped out of the study. Of those, 118 lacked SN samples, 2 sample size less than 5. The bioinformatic analysis pipelines were shown in Figure.1B and sFigure 1. The average age at onset of PD was 64-75 years old. After application of COMBAT, the batch effects were eliminated, and the data showed consistency (sFigure 2). To identify influential key genes in the SNc transcriptomic datasets of PD patients, the global Multiscale Embedded Gene Co-Expression Network Analysis (MEGENA) was utilized. A total of 82 parent-child gene network modules with module size >50 was identified from the SNc transcriptomic datasets of PD patients (Figure 1C). We found that PRKAR2B gene, which encodes PKA-RIIβ subunit, was one of the key hub genes among all functional genes in the module 50 (M50). Gene Ontology (GO) analysis demonstrated that differentially expressed genes (DEGs) in the M50 was closely linked to PD-related dopamine biosynthesis pathway (Figure 1C and D). In the M50, PRKAR2B gene was a top hub gene, and was associated with dopamine function-related genes, including DOPA decarboxylase (DDC), monoamine transporter (VMAT2) and dopamine transporter (DAT) (Figure 1E). Further, by using PPI network and CytoHubba plugin of Cytoscape, we observed that PRKAR2B was associated with dopamine metabolism, dopaminergic neurogenesis and PKA signaling pathway (Figure 1F). PRKAR2B was ranked top in hub genes based on the maximal clique centrality (MCC) algorithm in CytoHubba (Figure 1G). Moreover, to explore whether the PKA-RIIβ subunit in the SNc is involved in PD pathogenesis, we performed a correlation analysis by using SNc transcriptome sequencing datasets of PD patients. This analysis revealed a close correlation between PRKAR2B and other key genes related to dopamine metabolic pathways such as tyrosine hydroxylase (TH), DAT, DDC and VMAT2; oxidative stress pathways such as superoxide dismutase 1 (SOD1) and NADH: Ubiquinone Oxidoreductase Core Subunit S1 (NDUFS1); and apoptosis pathway such as BCL2 Associated X (BAX), BCL2 Antagonist/Killer 1 (BAK1) and Caspase 6 (CASP6) (Figure 1H). Notably, the meta-analysis showed that the gene expression of PRKAR2B was markedly decreased in human SNc of PD patients, as compared with non-PD individuals (Figure 1I). Taken together, these results unveil that PKA-RIIβ may be a key factor involved in dopamine synthesis, oxidative stress and apoptosis; and the lowered level of PKA-RIIβ may be linked to PD pathogenesis.

**Figure 1:**
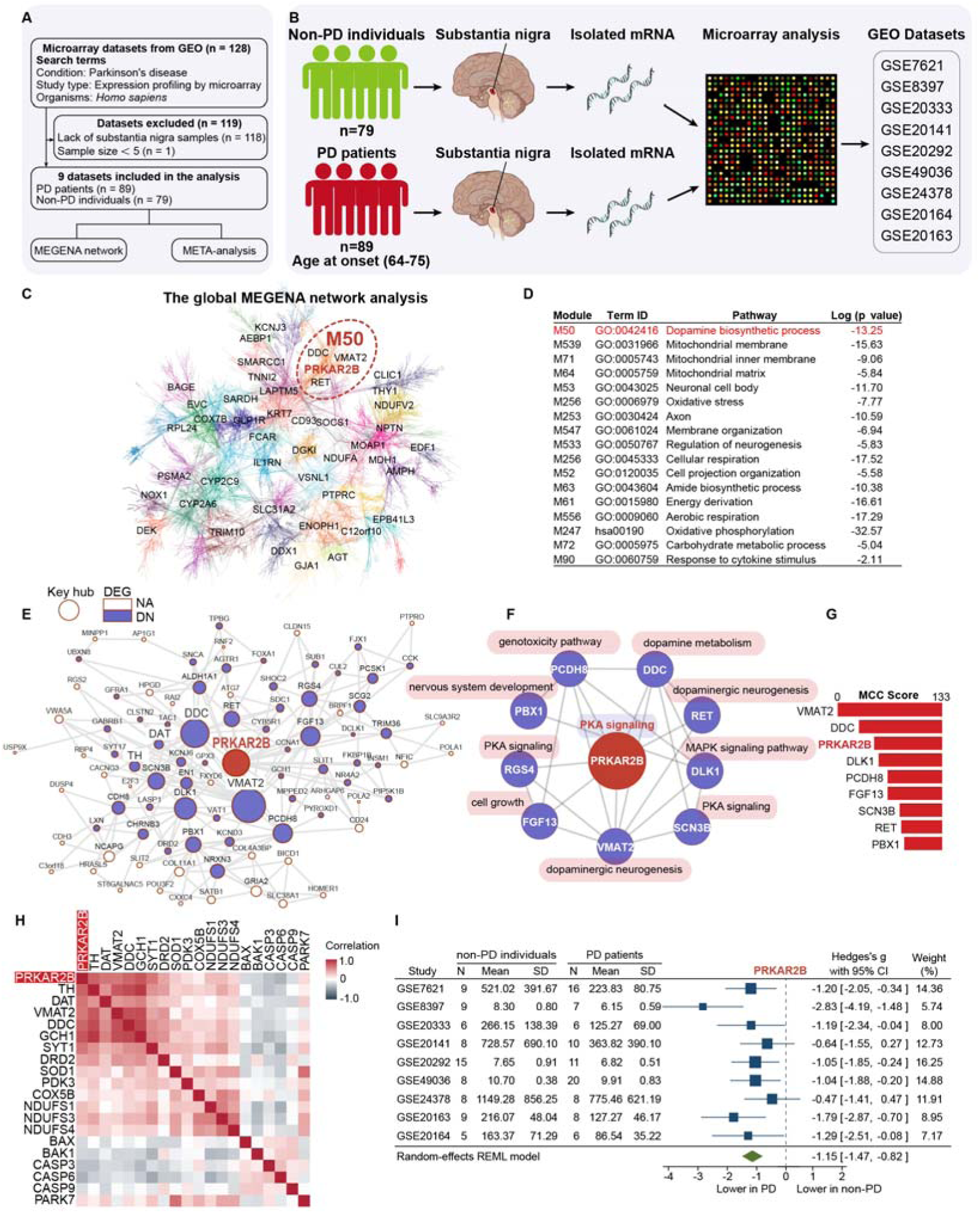
Human transcriptomic analysis shows that PKA-RIIβ may be linked to PD pathogenesis. **(A)** Enrollment of datasets in the analysis. **(B)** Schematic representation of dataset collection. Microarray-based gene expression datasets of postmortem substantia nigra from PD patients and non-PD individuals were collected from the GEO database. PD, Parkinson’s disease; GEO, gene expression omnibus. **(C)** The global MEGENA network of the SNc samples from PD patients. The modules at one particular compactness scale are represented by different colors. Highly connected hub genes are labeled with respective gene symbols. **(D)** GO and KEGG pathway enrichment analysis for genes in modules. The pathway with the smallest log *p* per module was shown in the table. GO, gene ontology; KEGG, kyoto encyclopedia of genes and genomes. **(E)** The top-ranked module M50 is highly enriched for the downregulated DEGs in PD. Key regulators are highlighted in burgundy circles. The size of a node is proportional to the node connectivity in the MEGENA network. **(F)** Hub genes in the module M50 were analyzed by CytoHubba Cytoscape plugin. **(G)** Hub genes in the module M50 were ranked by Maximal Clique Centrality (MCC) score. **(H)** Gene-gene correlation heatmap between PRKAR2B and genes of dopamine metabolism, oxidative stress and apoptosis in the SN of PD patients. **(I)** The meta-analysis of PRKAR2B in the SN between the postmortem PD patients and non-PD individuals, a random effect model was applied. All data are present as Hedges’s g with 95 % CI.

### RIIβ-KO mice show age-related spontaneous parkinsonism

Human PRKAR2B gene shares approximately 97% identity of nucleotide sequence with mouse PRKAR2B, showing a high level of sequence conservation (Figure 2A). In-situ hybridization data from the Allen Mouse Brain Atlas show a marked PRKAR2B expression in the SNc (Figure 2B). Consistent with the in situ data, a coexistence between RIIβ protein and TH was observed in the SNc dopaminergic neurons (TH^+^ neurons), validating that RIIβ is expressed in the SNc dopaminergic neurons (Figure 2C).

**Figure 2:**
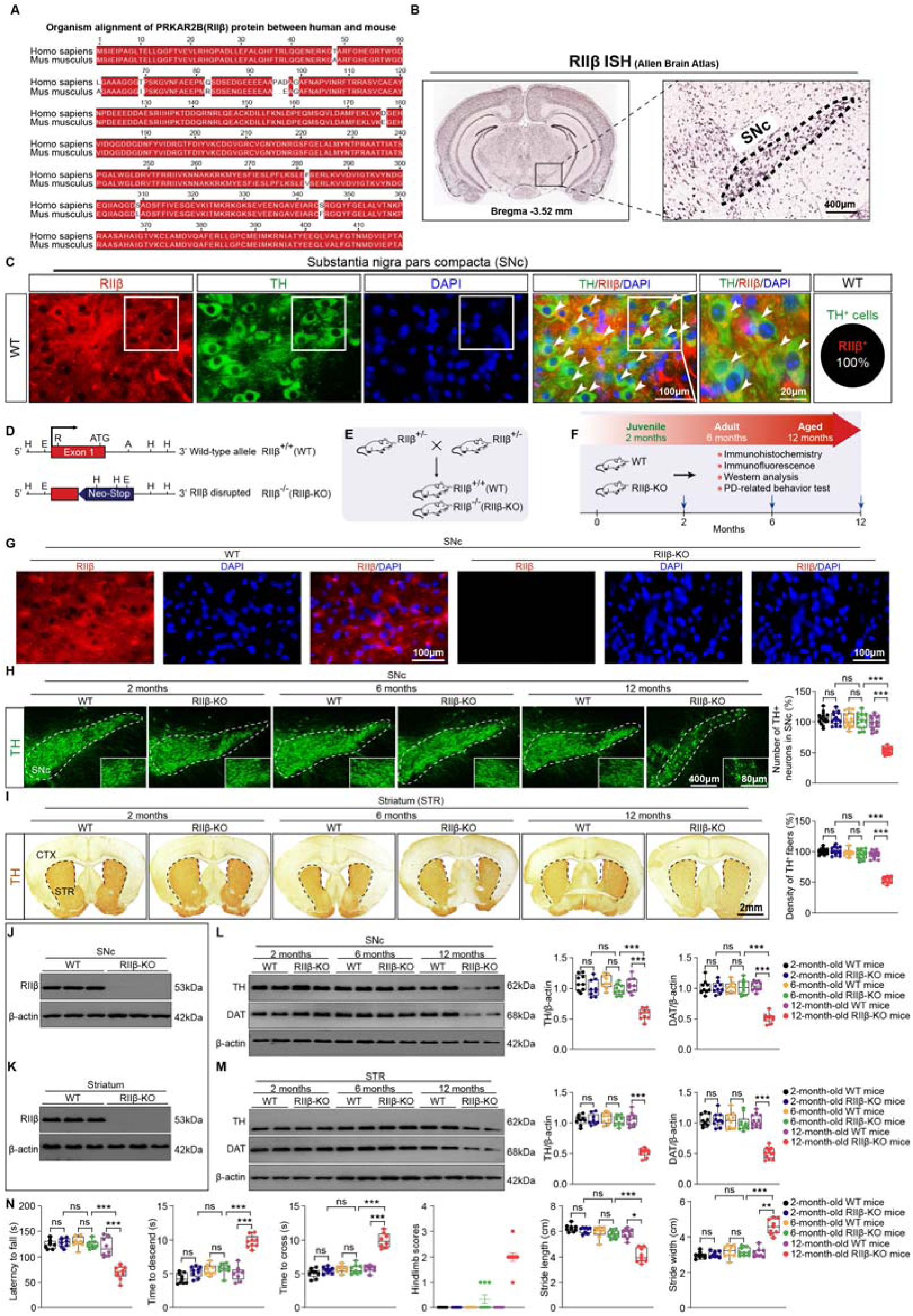
RIIβ-KO mice show an age-related spontaneous parkinsonism. **(A)** Multiple sequence alignment of the RIIβ protein sequence between Homo sapiens and Mus musculus. Identical residues are colored red. The alignment was done using ESPript 3.0 multiple sequence alignment program (https://espript.ibcp.fr/ESPript/ESPript/index.php). **(B)** In situ hybridization image from Allen Mouse Brain Atlas showed the expression pattern of RIIβ in the substantia nigra. SNc, substantia nigra pars compacta. **(C)** Representative double immunostaining of RIIβ with TH in SNc of WT mice. White arrow head indicates co-localization. Percentage of double-positive for TH (green) and RIIβ (red) was shown in the right. Scale bar, 100 μm for low-magnification images and 20 μm for high-magnification images, respectively. **(D)** Schematic strategy for generation of RIIβ knockout (RIIβ-KO) mice. The targeting vector substitutes the coding region of exon 1 of the PRKAR2B (RIIβ gene) with a neomycin resistance cassette (Neo). The indicated restriction enzyme sites are AatII (A), EcoRI (E), HindIII (H), and RsrI (R). **(E)** Breeding strategy for generation of RIIβ^+/+^(WT) mice and RIIβ^−/−^ (RIIβ-KO) mice. **(F)** Schematic representation of the experimental workflow. **(G)** Representative immunostaining of RIIβ in SNc of WT and RIIβ^−/−^ mice. Scale bar, 100 μm. **(H)** Representative images of TH and unbiased stereological counts of TH-positive neurons in SNc. Data are mean ± s.e.m.; n = 15 biologically independent animals; The two-way ANOVA was used for statistical analysis followed by Tukey’s multiple comparisons test. \*\*\**p* < 0.001. ns, not significant. Scale bars, 400 μm for low-magnification images and 80 μm for high-magnification images, respectively. **(I)** Representative images of TH staining and density of TH^+^ fibers in striatum. Data are mean ± s.e.m.; n = 15 biologically independent animals; The two-way ANOVA was used for statistical analysis followed by Tukey’s multiple comparisons test. \*\*\**p* < 0.001. ns, not significant. scale bar, 2 mm. **(J)** Representative immunoblots of RIIβ and β-actin in SNc of WT and RIIβ-KO mice (cropped blot images are shown, see sFigure 8 for full immunoblots). **(K)** Representative immunoblots of RIIβ and β-actin in striatum of WT and RIIβ-KO mice (cropped blot images are shown, see sFigure 8 for full immunoblots). **(L)** Representative immunoblots of TH, DAT, and β-actin (cropped blot images are shown, see sFigure 8 for full immunoblots) and quantification of TH and DAT levels in SNc. Data are mean ± s.e.m.; n = 9 biologically independent animals. The two-way ANOVA was used for statistical analysis followed by Tukey’s multiple comparisons test. \*\*\**p* < 0.001. ns, not significant. **(M)** Representative immunoblots of TH, DAT, and β-actin (cropped blot images are shown, see sFigure 8 for full immunoblots) and quantification of TH and DAT levels in striatum. Data are mean ± s.e.m.; n = 9 biologically independent animals. The two-way ANOVA was used for statistical analysis followed by Tukey’s multiple comparisons test. \*\*\**p* < 0.001. ns, not significant. **(N)** Fall latency from an accelerating rotarod, time to traverse beam apparatus, time to descend pole, hindlimb clasping reflex score and gait analysis. Data are mean ± s.e.m.; n = 9 biologically independent animals. The two-way ANOVA was used for statistical analysis followed by Tukey’s multiple comparisons test. \**p* < 0.05, \*\**p* < 0.01 and \*\*\**p* < 0.001. ns, not significant.

To explore whether PKA-RIIβ is linked to PD pathogenesis, PKA-RIIβ gene knockout (RIIβ-KO) mice were employed. The strategy for generating RIIβ-KO mice; and the strategy for breeding RIIβ^−/−^ mice (RIIβ-KO) mice were illustrated in Figure 2D and E. By using molecular and behavioral metrics of PD, we assessed the association of PKA-RIIβ with PD; the experimental strategy was shown in Figure 2F. We found that RIIβ subunit was absent in the SNc and the striatum (STR) of RIIβ-KO mice (Figure2G, J and K, sFigure 3A and B). Notably, at 12 months of age, RIIβ-KO mice showed a loss of dopaminergic neurons (∼50% loss) in the SNc (Figure 2H) and a decrease in TH^+^ fibers density in the STR (∼50%), as compared with that of WT mice (Figure 2I); whereas no difference was observed at 3 months and 6 months of age. The protein levels of both TH and DAT were reduced in the SNc and the STR of the RIIβ-KO mice at 12 months of age, but not at 3 months and 6 months of age (Figure 2L and M). Pole descent, rotarod test, beam traversal, hindlimb clasping reflexes and gait test showed that motor coordination and balance were impaired in 12-month-old RIIβ-KO mice (Figure 2N). Taken together, these results reveal that the RIIβ is expressed in the SNc dopaminergic neurons. RIIβ gene knockout induces spontaneous parkinsonism in aged mice, suggesting that RIIβ deficiency may be associated with PD pathogenesis. The age-related spontaneous parkinsonism of RIIβ-KO mice may also echo the lowered level of RIIβ in the SNc of PD patients.

### RIIβ reexpression in SNc dopaminergic neurons relieves spontaneous parkinsonism of RII**β**-KO mice

The global knockout of the RΙΙβ gene leads to an age-related spontaneous parkinsonism; the RΙΙβ subunit is expressed in both SNc and STR. Thus, we asked where the RIIβ is involved in PD pathogenesis. As described previously, mice were engineered to reexpress RIIβ in response to Cre recombinase activity^49^. A loxP-flanked cassette containing the neomycin resistance gene was inserted between the transcription start site and the ATG codon of the RIIβ gene (Figure 3A). The neo-STOP sequence is deleted when Cre-recombinase is expressed in RΙΙβ^lox/−^mice, and thus a loxP locus of the RIIβ gene is eventually left behind (Figure 3A). Cre recombinase transgenes may become active that may result in germline recombination in progeny^57^. Thus, for avoiding germline recombination, we bred the DAT-cre transgene onto the RIIβ^−/−^ background and then mated them to RIIβ^lox/lox^mice to obtain DAT-Cre/RIIβ^lox/−^ mice (Figure. 3B). By adopting same breeding strategy, D1R-cre transgene or A2A-cre transgene were also used to reexpress RIIβ in dopamine D1 receptor (D1R)-expressing striatal medium-sized spiny neurons (MSNs) or dopamine D2 receptor (D2R)-expressing MSNs, respectively (Figure 3C and D). In DAT-Cre/RIIβ^lox/−^ mice, RΙΙβ was specifically reexpressed in the dopaminergic neurons of the SNc (Figure 3F and G). In D1R-Cre/RΙΙβ^lox/−^ mice, RΙΙβ was reexpressed in D1R striatal MSNs; and in A2A-Cre/RΙΙβ^lox/−^ mice, RΙΙβ was reexpressed in D2R striatal MSNs (sFigure 4A and B). Further, we evaluated the parkinsonism of these mice; and the experimental strategy was illustrated in Figure 3E. We found that the loss of SNc dopaminergic neurons (89 % versus 46 %) and the decreased TH^+^ fibers density (92 % versus 50 %) was relieved in DAT-Cre/RIIβ^lox/−^ mice, but not in D1R-Cre/RΙΙβ^lox/−^ mice and A2A-Cre/RΙΙβ^lox/−^ mice (Figure 3H and I). The decreased protein level of TH and DAT in the SNc and the STR were normalized in DAT-Cre/RIIβ^lox/−^ mice (Figure 3J and K). The parkinsonism was mitigated in DAT-Cre/RIIβ^lox/−^ mice (Figure 3L). Together, these observations demonstrate that RIIβ reexpression in the SNc dopaminergic neurons relieves the spontaneous parkinsonism induced by RΙΙβ deficiency, unveiling that RIIβ in the SNc dopaminergic neurons may underlie PD pathogenesis.

**Figure 3:**
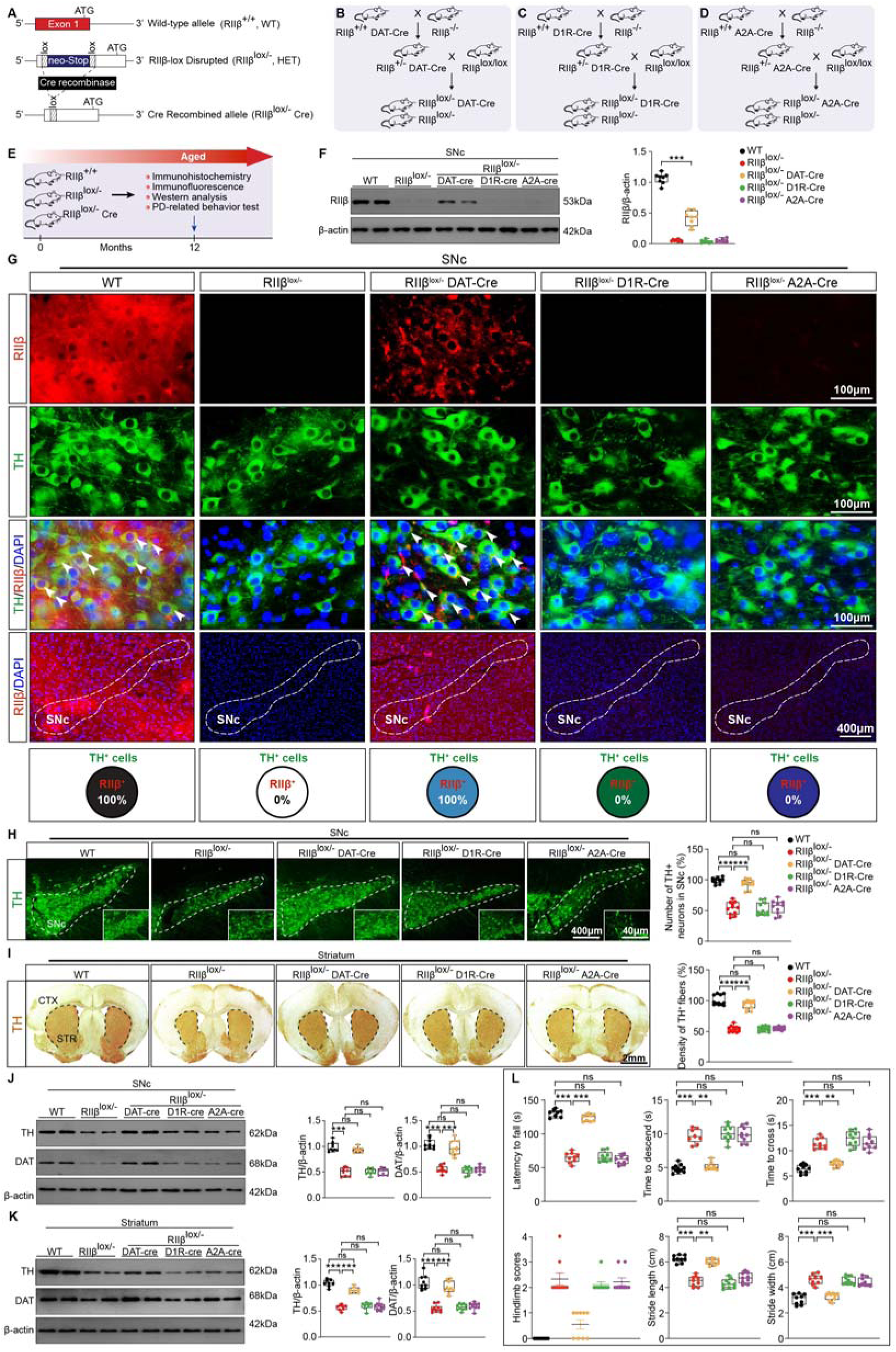
RIIβ reexpression in SNc dopaminergic neurons relieves spontaneous parkinsonism of RIIβ-KO mice. **(A)** Strategy for generation of RIIβ^lox/−^and RIIβ^lox/−^/Cre mice. **(B)** Breeding strategy for generation of DAT-Cre mice with dopaminergic neuron-specific RIIβ reexpression (RIIβ^lox/−^ DAT-Cre). **(C)** Breeding strategy for generation of D1R-Cre mice with dopamine receptor 1-expressing medium spiny neuron-specific RIIβ reexpression (RIIβ^lox/−^ D1R-Cre). **(D)** Breeding strategy for generation of A2A-Cre mice with striatal adenosine A2A receptor neuron-specific RIIβ reexpression (RIIβ^lox/−^A2A-Cre). **(E)** Schematic representation of the experimental workflow. **(F)** Representative immunoblots of RIIβ and β-actin (cropped blot images are shown, see sFigure 9 for full immunoblots) and quantification of RIIβ levels in SNc. Data are mean ± s.e.m.; n = 9 biologically independent animals. The two-way ANOVA was used for statistical analysis followed by Tukey’s multiple comparisons test. \*\*\**p* < 0.001. **(G)** Representative double immunostaining of RIIβ with TH in SNc of WT, RIIβ^lox/−^, RIIβ^lox/−^ DAT-Cre, RIIβ^lox/−^ D1R-Cre and RIIβ^lox/−^ A2A-Cre mice. Scale bar, 400 μm for low-magnification images and 100 μm for high-magnification images, respectively. **(H)** Representative images of TH and unbiased stereological counts of TH-positive neurons in SNc. Data are mean ± s.e.m.; n = 11 biologically independent animals; The two-way ANOVA was used for statistical analysis followed by Tukey’s multiple comparisons test. \*\*\**p* < 0.001. ns, not significant. Scale bars, 400 μm for low-magnification images and 40 μm for high-magnification images, respectively. **(I)** Representative images of TH staining and density of TH^+^ fibers in striatum. Data are mean ± s.e.m.; n = 11 biologically independent animals; The two-way ANOVA was used for statistical analysis followed by Tukey’s multiple comparisons test. \*\*\**p* < 0.001. ns, not significant. Scale bar, 2 mm. **(J)** Representative immunoblots of TH, DAT, and β-actin (cropped blot images are shown, see sFigure 9 for full immunoblots) and quantification of TH and DAT levels in SNc. Data are mean ± s.e.m.; n = 9 biologically independent animals. The two-way ANOVA was used for statistical analysis followed by Tukey’s multiple comparisons test. \*\*\**p* < 0.001. ns, not significant. **(K)** Representative immunoblots of TH, DAT, and β-actin (cropped blot images are shown, see sFigure 9 for full immunoblots) and quantification of TH and DAT levels in STR. Data are mean ± s.e.m.; n = 9 biologically independent animals. The two-way ANOVA was used for statistical analysis followed by Tukey’s multiple comparisons test. \*\*\**p* < 0.001. ns, not significant. **(L)** Fall latency from an accelerating rotarod, time to traverse beam apparatus, time to descend pole, hindlimb clasping reflex score and gait analysis. Data are mean ± s.e.m.; n = 9 biologically independent animals. The two-way ANOVA was used for statistical analysis followed by Tukey’s multiple comparisons test. \*\**p* < 0.01 and \*\*\**p* < 0.001. ns, not significant.

**Figure 4:**
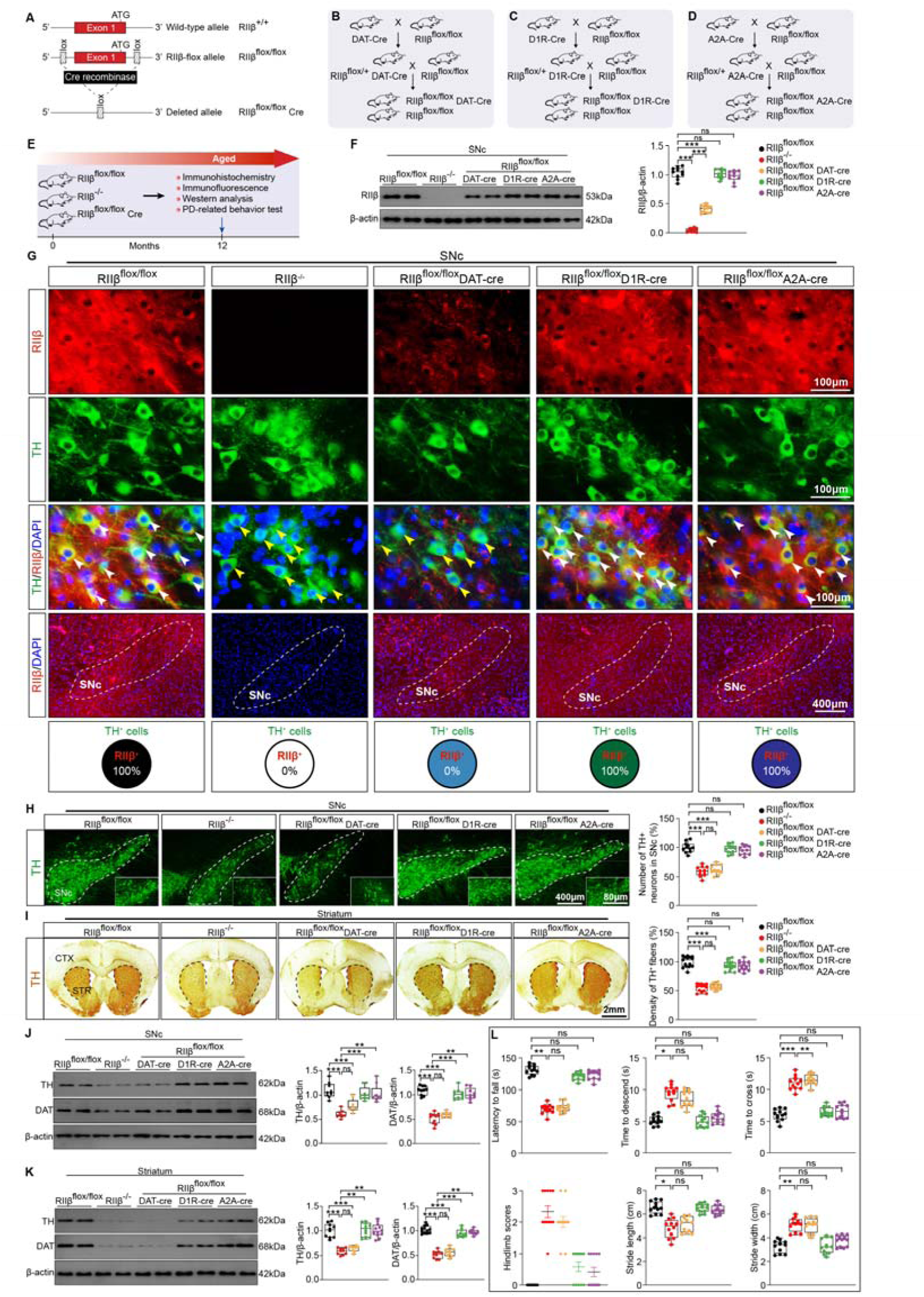
RIIβ deficiency in SNc dopaminergic neurons induces spontaneous parkinsonism. **(A)** Strategy for generation of RIIβ^flox/flox^ and RIIβ^flox/flox^ Cre mice. **(B)** Breeding strategy for generation of DAT-Cre mice with SNc dopaminergic neuron-specific RIIβ knockout (RIIβ^flox/flox^ DAT-Cre). **(C)** Breeding strategy for generation of D1R-Cre mice with dopamine receptor 1-expressing medium spiny neuron-specific RIIβ knockout (RIIβ^flox/flox^ D1R-Cre). **(D)** Breeding strategy for generation of A2A-Cre mice with striatal adenosine A2A receptor neuron-specific RIIβ knockout (RIIβ^flox/flox^A2A-Cre). **(E)** Schematic representation of the experimental workflow. **(F)** Representative immunoblots of RIIβ and β-actin (cropped blot images are shown, see sFigure 9 for full immunoblots) and quantification of RIIβ levels in SNc. Data are mean ± s.e.m.; n = 9 biologically independent animals. The two-way ANOVA was used for statistical analysis followed by Tukey’s multiple comparisons test. \*\*\**p* < 0.001. ns, not significant. **(G)** Representative immunostaining of RIIβ with TH in SNc of RIIβ^flox/flox^, RIIβ^−/−^, RIIβ^flox/flox^ DAT-cre, RIIβ^flox/flox^ D1R-cre and RIIβ^flox/flox^ A2A-cre mice. White arrow head indicates co-localization, yellow arrow head indicates TH-positive neurons without RIIβ. 400 μm for low-magnification images and 100 μm for high-magnification images, respectively. **(H)** Representative images of TH and unbiased stereological counts of TH-positive neurons in SNc. Data are mean ± s.e.m.; n = 10 biologically independent animals; The two-way ANOVA was used for statistical analysis followed by Tukey’s multiple comparisons test. \*\*\**p* < 0.001. ns, not significant. Scale bars, 400 μm for low-magnification images and 80 μm for high-magnification images, respectively. **(I)** Representative images of TH staining and density of TH^+^ fibers in striatum. Data are mean ± s.e.m.; n = 12 biologically independent animals; The two-way ANOVA was used for statistical analysis followed by Tukey’s multiple comparisons test. \*\*\**p* < 0.001. ns, not significant. Scale bar, 2 mm. **(J)** Representative immunoblots of TH, DAT, and β-actin (cropped blot images are shown, see sFigure 9 for full immunoblots) and quantification of TH and DAT levels in SNc. Data are mean ± s.e.m.; n = 9 biologically independent animals. The two-way ANOVA was used for statistical analysis followed by Tukey’s multiple comparisons test. \*\**p* < 0.01 and \*\*\**p* < 0.001. ns, not significant. **(K)** Representative immunoblots of TH, DAT, and β-actin (cropped blot images are shown, see sFigure 9 for full immunoblots) and quantification of TH and DAT levels in STR. Data are mean ± s.e.m.; n = 9 biologically independent animals. The two-way ANOVA was used for statistical analysis followed by Tukey’s multiple comparisons test. \*\**p* < 0.01 and \*\*\**p* < 0.001. ns, not significant. **(L)** Fall latency from an accelerating rotarod, time to traverse beam apparatus, time to descend pole, hindlimb clasping reflex score and gait analysis. Data are mean ± s.e.m.; n = 9 biologically independent animals. The two-way ANOVA was used for statistical analysis followed by Tukey’s multiple comparisons test. \**p* < 0.05, \*\**p* < 0.01 and \*\*\**p* < 0.001. ns, not significant.

### RIIβ deficiency in SNc dopaminergic neurons causes spontaneous parkinsonism

To further validate whether RΙΙβ deficiency in the SNc dopaminergic neurons may cause PD pathogenesis, we used the Cre-loxP system to generate cell type-specific RΙΙβ knockout mice. Mice were engineered to knockout RIIβ gene in response to Cre recombinase activity (Figure 4A). The SNc dopaminergic neuron-specific RIIβ-deficient mice (DAT-Cre/RIIβ^flox/flox^), striatal D1R MSNs-specific RIIβ-deficient mice (D1R-Cre/RIIβ^flox/flox^) and striatal D2R MSNs-specific RIIβ-deficient mice (A2A-Cre/RIIβ^flox/flox^) were generated using the strategy described in Figure 4B-D. We assessed the spontaneous parkinsonism of these mice at 12 months of age; the experimental procedure was described in Figure 4E. We observed that RIIβ level was specifically reduced in SNc of DAT-Cre/RIIβ^flox/flox^mice, but not in the SNc of D1R-Cre/RΙΙβ^flox/flox^ mice and A2A-Cre/RΙΙβ^flox/flox^ mice (Figure 4F and G). The RIIβ expression levels were reduced in STR of D1R-Cre/RΙΙβ^flox/flox^ mice and A2A-Cre/RΙΙβ^flox/flox^mice (sFigure 5A and B). A loss of dopaminergic neurons (∼40 % loss) and a decrease in TH^+^ fibers density (∼45 %) were observed in DAT-Cre/RIIβ^flox/flox^ mice, as compared with that of WT mice, at 12 months of age; but were not in D1R-Cre/RΙΙβ^flox/flox^ mice or A2A-Cre/RΙΙβ^flox/flox^ mice (Figure 4 H and I). TH and DAT immunoreactivity was reduced in SNc and STR of 12-month-old DAT-Cre/RIIβ^flox/flox^ mice, but not in D1R-Cre/RΙΙβ^flox/flox^ mice or A2A-Cre/RΙΙβ^flox/flox^ mice (Figure 4 J and K). Notably, the motor coordination and balance were impaired in the 12-month-old DAT-Cre/RIIβ^flox/flox^mice, but not in D1R-Cre/RΙΙβ^flox/flox^mice or A2A-Cre/RΙΙβ^flox/flox^mice (Figure 4 L). Together, these results show that the spontaneous parkinsonism can be developed in 12-month-old DAT-Cre/RIIβ^flox/flox^ mice, highlighting that RIIβ deficiency in SNc dopaminergic neurons may lead to PD.

### Decreased PKA activity and reduced dopamine synthesis in SNc dopaminergic neurons of RIIβ-KO mice

To explore the mechanisms by which RIIβ-PKA in SNc dopaminergic neurons underlie PD pathogenesis, single-nucleus RNA-sequencing (snRNA-seq) and whole-genome RNA-sequencing (RNA-Seq) analyses were employed. We microdissected the SNc from fresh brain slices for RNA-Seq, and processed single-nucleus suspensions through the 10× Genomics Chromium Controller as previously described^58^. The microdissection was mapped to ensure accuracy and reproducibility; and the analysis strategy was illustrated in Figure 5A. Our initial database included individual transcriptomic profiles from RIIβ-KO (n= 11030 cells) and WT (n = 7118 cells) samples. According to the reported markers, we assigned single cells to a given cell type, including astrocytes (AQP4, ALDH1L1, AGT, GFAP), endothelial cells (FLT1), microglia (CTSS, CSF1R, CX3CR1), neurons (SNAP25, SYP, SYT1, ENO2), oligodendrocytes (MAG, MOG, MBP, MOBP), oligodendrocyte precursor cells (CSPG4, VCAN, PDGFRA) and pericytes (VTN), enabling automated annotation of the cell types in SNc (Figure 5B). We then proceeded to apply unsupervised, iterative clustering to distinguish molecularly distinct neuronal clusters. We identified three different SNc neuronal clusters according to expression of the genes for the synthesis and packaging of dopamine, GABA and glutamate. In particular, the genes of TH and DAT (SLC6A3^+^) were expressed in the neuronal clusters of 2,5,8 and 11. SLC17A6, which encodes vesicular glutamate transporter 2, was expressed in the neuronal clusters of 3,4,10 and 13. SLC32A1, which encodes vesicular GABA transporter (VGAT), was expressed in the neuronal clusters of 0,1,6,7,9,12 and 13 (Figure 5 C and D). Thus, we nominally categorized SLC6A3^+^ clusters as dopaminergic, SLC32A1^+^ clusters as GABAergic and SLC17A6^+^ clusters as glutamatergic neurons (Figure 5 E).

**Figure 5:**
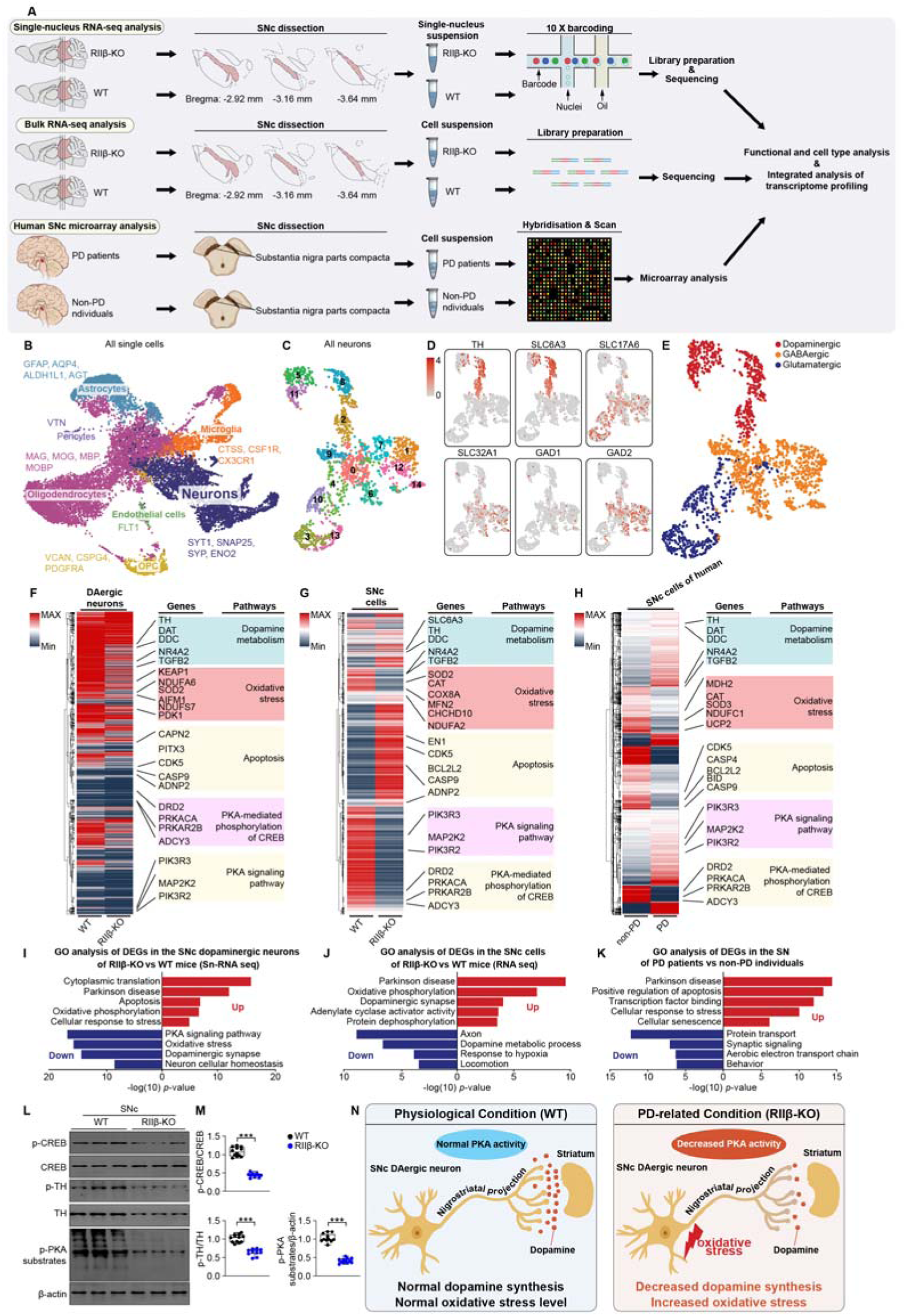
Decreased PKA activity and reduced dopamine synthesis in SNc dopaminergic neurons of RIIβ-KO mice. **(A)** Schematic representation of the experimental workflow. sn-RNAseq: WT n = 5; RIIβ-KO n = 5. Bulk RNA-seq: WT n = 3; RIIβ-KO n = 3. **(B)** UMAP visualization of seven transcriptionally distinct clusters expressing canonical markers. Neurons (SNAP25, SYP, SYT1, ENO2); Astrocytes (AQP4, ALDH1L1); Oligodendrocytes (MAG, MOG, MBP, MOBP); OPC, oligodendrocyte precursor cells (CSPG4, VCAN, PDGFRA); Microglia (CTSS, CSF1R, CX3XR1); Pericytes (VTN); Endothelial cells (FLT1). (n = 18148 cells). **(C)** UMAP visualization of 14 transcriptionally distinct clusters of neurons after the first iteration of unsupervised clustering. **(D)** Normalized expression of TH, SLC6A3, GAD1, GAD2, SLC32A1, and SLC17A6 in each cell shown on UMAP plot after the second iteration of unsupervised clustering of neurons. **(E)** Neurons were classified as dopaminergic, GABAergic or glutamatergic based on the expression of canonical markers (n = 1750 neurons). **(F)** Hierarchical clustered heatmap of gene expression profiles at sn-RNA seq resolution in the SNc dopaminergic neurons of WT and RIIβ-KO mouse). **(G)** Hierarchical clustered heatmap of gene expression profiles at bulk RNA-seq resolution in the SNc of WT and RIIβ-KO mouse. **(H)** Hierarchical clustered heatmap of gene expression profiles in the SN of humans (PD patients and non-PD individuals). **(I)** GO analysis of DEGs in the SNc dopaminergic neurons of WT and RIIβ-KO mouse (Sn-RNA seq). **(J)** GO analysis of DEGs in the SNc cells of WT and RIIβ-KO mouse (RNA seq). **(K)** GO analysis of DEGs in the SN of PD patients and non-PD individuals. **(L)** Representative immunoblots of p-CREB, CREB, p-PKA substrates and β-actin in SNc of WT and RIIβ^−/−^ mice (cropped blot images are shown, see sFigure 9 for full immunoblots). **(M)** Quantification of p-CREB, p-TH and p-PKA substrates levels. Data are mean ± s.e.m.; n = 12 biologically independent animals. Unpaired two tailed Student’s t-tests were used for statistical analyses. \*\*\**p* < 0.001. **(N)** Diagram of the mechanisms underlying the dopamine metabolism dysfunction of SNc dopaminergic neurons in RIIβ-KO mice.

To analyze the mechanistic changes in SNc dopaminergic neurons, differentially expressed genes (DEGs) analysis was performed. DEGs associated with dopamine metabolism (7 genes), PKA activity (8 genes), oxidative stress (7 genes) and apoptosis (7 genes) were observed (Figure 5F and G, sFigure 6A). Notably, these DEGs-related pathways were also observed in the SNc of PD patients (Figure 5H, sFigure 6B). GO and KEGG analyses showed that the pathways associated with PD, apoptosis and oxidative stress were induced, while the pathways associated with dopamine metabolic process, dopaminergic synapse and PKA signaling pathway were inhibited in RIIβ-KO mice (Figure 5I and J). In the SNc of PD patients, GO and KEGG analyses of DEGs also showed that pathways associated with PD and apoptosis were induced, whereas aerobic electron transport chain and synaptic signaling were inhibited (Figure 5K). Of note, PKA activity was decreased in SNc dopaminergic neurons of RIIβ-KO mice, as indicated by decreased phosphorylated protein levels of CREB and PKA substrates (Figure 5L and M). TH phosphorylation at Ser 40 was reduced in the SNc of RIIβ-KO mice (Figure 5 L and M). Together, these results indicate that lowered level or deficiency of RIIβ may lead to a decreased PKA activity, causing a reduced dopamine synthesis in SNc dopaminergic neurons, which may be a key mechanism of PD pathogenesis (Figure 5N).

### Conversion of Type-II PKA to Type-I PKA in SNc dopaminergic neurons underlies PD pathogenesis of RIIβ-KO mice

To analyze the expression pattern of PKA subunits in the SNc dopaminergic neurons of RIIβ-KO mice, single-nucleus sequencing and UMAP’s dimensionality reduction algorithm in conjunction with unsupervised cell clustering was utilized. We identified six types of SNc dopaminergic neurons (Figure 6A). The transcriptional alterations of PKA subunits were analyzed between RIIβ-KO mice and WT mice. RIIβ deficiency resulted in a PRKAR1A increase of approximately 30 %; whereas, levels of PRKAR1B, PRKAR2A and PRKARCA were reduced, in SNc dopaminergic neurons of RIIβ-KO mice, as compared with WT controls (Figure 6B). These findings showed that there was a conversion of Type-II PKA to Type-I PKA, revealing a decrease in C subunits (approximately 20 % loss of C subunits) and R subunits (approximately 60% loss of total R subunits), and reduced total PKA activity in SNc dopaminergic neurons of RIIβ-KO mice (Figure 6B). The functional enrichment analysis of DEGs between RIIβ-KO and WT mice showed a decreased PKA activity, decreased dopamine metabolism, and increased oxidative stress in the SNc dopaminergic neurons of RIIβ-KO mice (Figure 6C). Both up- and down-regulated DEGs were closely related to PKA activity (9 genes), dopamine metabolic process (5 genes), oxidative stress (8 genes) and apoptosis (6 genes) (Figure 6D). Correlation analysis showed a close correlation among PKA activity, dopamine metabolic process, oxidative stress and neuronal apoptosis (Figure 6E). Together, these analyses uncovered that RIIβ deficiency leads to a conversion of Type-II PKA to Type-I PKA, showing decreased PKA activity, impaired dopamine synthesis, increased oxidative stress, and neuronal apoptosis in the SNc dopaminergic neurons (Figure 6F and G), suggesting that the conversion of PKA kinase system caused by RIIβ deficiency may account for the PD-related phenotypes.

**Figure 6:**
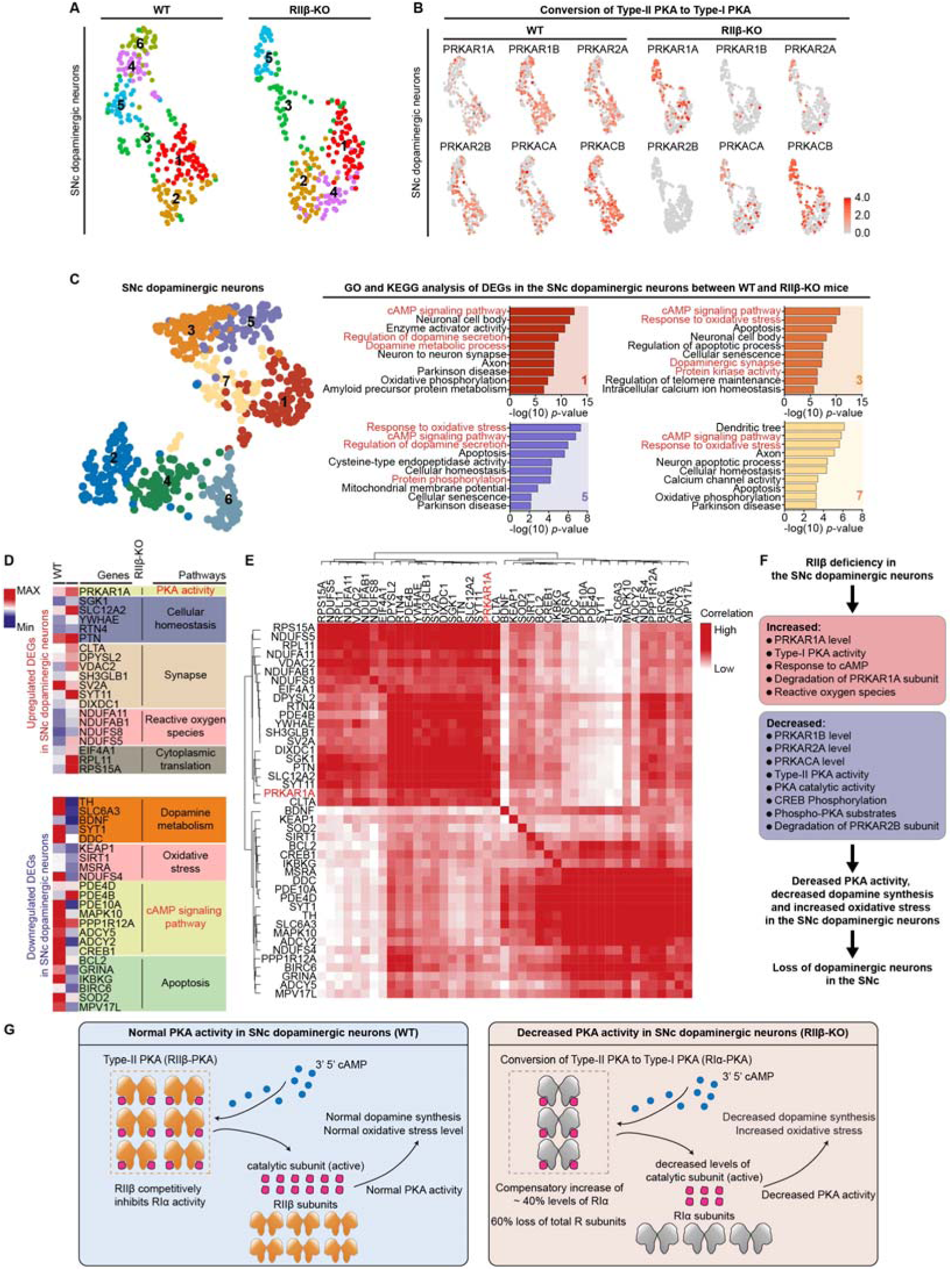
Conversion of Type-II PKA to Type-I PKA in SNc dopaminergic neurons may underlie PD pathogenesis of RIIβ-KO mice. **(A)** Unsupervised clustering of SNc dopaminergic neuronal types represented in UMAP plots. Cell-type clusters are color-coded. **(B)** Expression patterns of PKA subunits in SNc dopaminergic neurons of WT and RIIβ-KO mice. **(C)** GO and KEGG analysis of DEGs in SNc dopaminergic neurons between WT and RIIβ-KO mice. **(D)** Most significantly altered (based on *p* rank) genes (rows) in SNc dopaminergic neurons of RIIβ-KO and WT mice (columns). **(E)** Gene-gene correlation heatmap of the DEGs in SNc dopaminergic neurons between RIIβ-KO and WT mice. **(F)** Schematic diagram showing the mechanism of the decreased PKA activity in SNc dopaminergic neurons of RIIβ-KO mice. **(G)** Diagram of the mechanisms underlying the decreased PKA activity of SNc dopaminergic neurons in WT and RIIβ-KO mice.

### RIIβ deficiency aggravates MPTP-induced PD phenotypes

snRNA-seq and RNA-seq analyses revealed that RIIβ deficiency may cause increased oxidative stress. Meta-analysis of the SNc transcriptome sequencing datasets from MPTP-induced PD mice showed that RIIβ level was decreased in the SNc of MPTP-treated mice, suggesting that a lowered RIIβ level may be associated with a higher level of oxidative stress in the SNc of PD mice (Figure 7A). Thus, we test whether RIIβ deficiency may affect MPTP-induced PD phenotypes. The experimental strategy was illustrated in Figure 7B. In MPTP-treated mice, RIIβ deficiency caused a marked loss of dopaminergic neurons (49 % versus 33 %) (Figure 7C), a decrease in TH^+^ fibers density (Figure 7D), and a reduction in TH and DAT protein levels (Figure 7E and F), Pole descent, rotarod test, beam traversal, hindlimb clasping reflexes and gait test showed that RIIβ deficiency worsened the impaired motor function in MPTP-treated mice (Figure 7G). Together, these results suggest that RIIβ-PKA may be involved in the regulation of mitochondrial oxidation in SNc dopaminergic neurons; and lowered level or deficiency of RIIβ may be related to elevated oxidative stress, and thus underlie PD pathogenesis.

**Figure 7:**
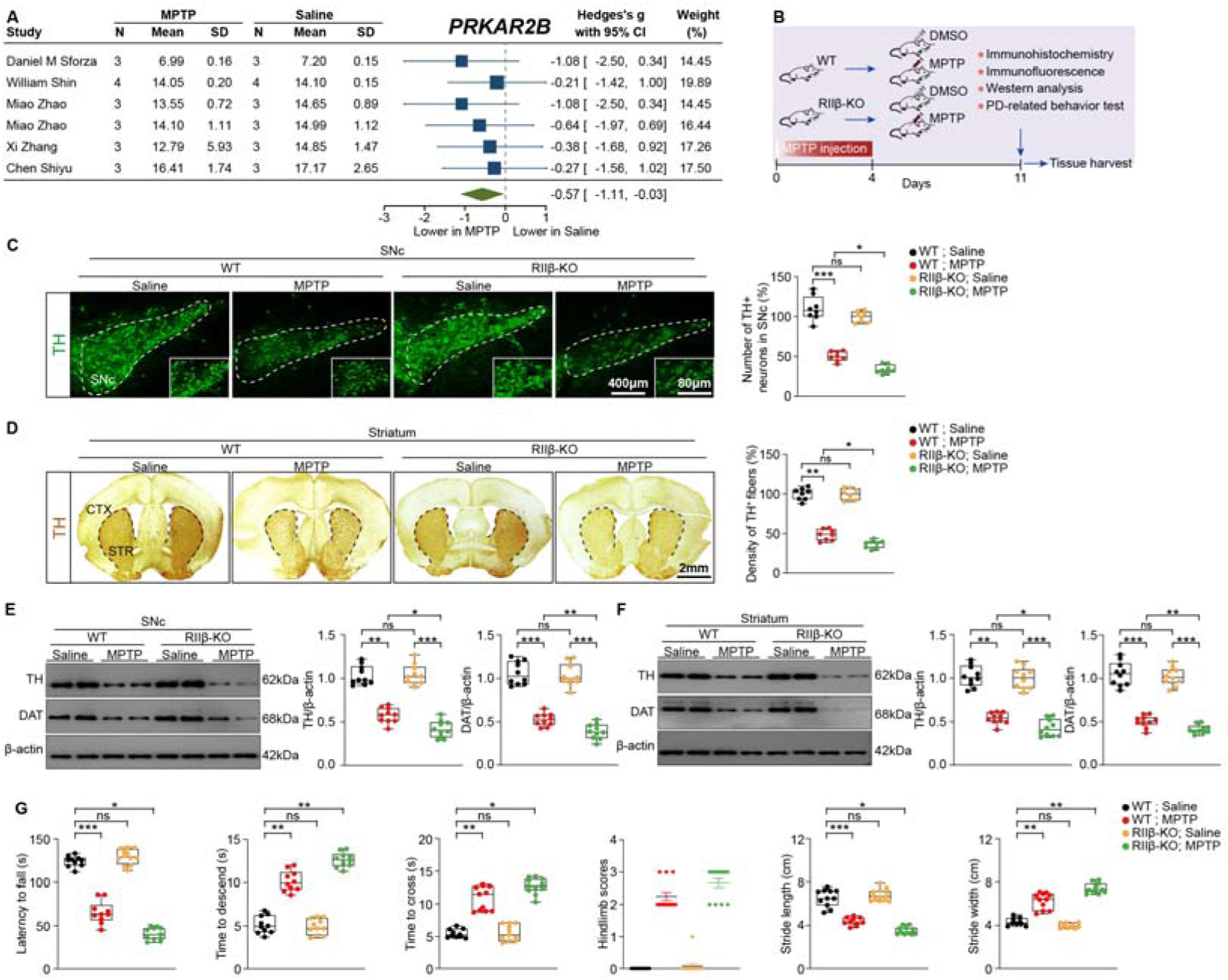
RIIβ deficiency aggravates MPTP-induced PD phenotypes. **(A)** The meta-analysis of PRKAR2B levels in the SNc between MPTP-induced PD mice and control mice, a random effect model was applied. All data are present as Hedges’s g with 95% CI. **(B)** Schematic representation of the experimental workflow. **(C)** Representative images of TH and unbiased stereological counts of TH-positive neurons in SNc. Data are mean ± s.e.m.; n = 11 biologically independent animals; The two-way ANOVA was used for statistical analysis followed by Tukey’s multiple comparisons test. \**p* < 0.05 and \*\*\**p* < 0.001. ns, not significant. Scale bars, 400 μm for low-magnification images and 80 μm for high-magnification images, respectively. **(D)** Representative images of TH staining and density of TH^+^ fibers in striatum. Data are mean ± s.e.m.; n = 11 biologically independent animals; The two-way ANOVA was used for statistical analysis followed by Tukey’s multiple comparisons test. \**p* < 0.05 and \*\**p* < 0.01. ns, not significant. Scale bar, 2 mm. **(E)** Representative immunoblots of TH, DAT, and β-actin in SNc (cropped blot images are shown, see sFigure 10 for full immunoblots) and quantification of TH and DAT levels. Data are mean ± s.e.m.; n = 9 biologically independent animals. The two-way ANOVA was used for statistical analysis followed by Tukey’s multiple comparisons test. \**p* < 0.05, \*\**p* < 0.01 and \*\*\**p* < 0.001. ns, not significant. **(F)** Representative immunoblots of TH, DAT, and β-actin (cropped blot images are shown, see sFigure 10 for full immunoblots) and quantification of TH and DAT levels in STR. Data are mean ± s.e.m.; n = 9 biologically independent animals. The two-way ANOVA was used for statistical analysis followed by Tukey’s multiple comparisons test. \**p* < 0.05, \*\**p* < 0.01 and \*\*\**p* < 0.001. ns, not significant. **(G)** Fall latency from an accelerating rotarod, time to traverse beam apparatus, time to descend pole, hindlimb clasping reflex score and gait analysis. Data are mean ± s.e.m.; n = 9 biologically independent animals. The two-way ANOVA was used for statistical analysis followed by Tukey’s multiple comparisons test. \**p* < 0.05, \*\**p* < 0.01 and \*\*\**p* < 0.001. ns, not significant.

### AAV-based gene therapy targeting PKA-RIIβ relieves MPTP-induced PD symptoms

To explore whether an increase in PKA activity induced by RIIβ mutation in the SNc dopaminergic neurons may treat PD, an adeno-associated virus (AAV)-based gene therapy was employed. MPTP treatment increases oxidative stress in the SNc dopaminergic neurons in mice^59^, and RIIβ-PKA may be involved in the oxidative process in the SNc dopaminergic neurons; thus, MPTP-induced PD mouse model was used. A mutation of serine 112 to alanine of RIIβ subunit (RIIβ-S112A), a P-site mutant that cannot be phosphorylated, can protect C subunits, and thus heighten PKA activity^25,60–63^(Figure 8A). The experimental strategy was shown in Figure 8B. We expressed RIIβ-S112A in dopaminergic neurons or all neurons in the SNc by bilateral stereotaxic injection of Flex-RIIβ-S112A or hsyn-RIIβ-S112A in DAT-cre mice; and GFP fluorescence was visualized (Figure 8C). We observed that the AAV-RIIβ-S112A was robustly expressed in the SNc in the third week after the injection (sFigure 7). The strategy for the study of gene therapy for PD was illustrated in Figure 8D. AAV-delivered Flex-RIIβ-S112A and hsyn-RIIβ-S112A in the SNc normalized the levels of p-CREB and p-PKA substrates, mitigated the loss of dopaminergic neurons, relieved the decrease of TH^+^ fibers density, rescued the protein levels of TH and DAT, and alleviated the impaired motor coordination and balance in MPTP-treated mice (Figure 8 E-J). Together, these findings demonstrate that enhancement of PKA activity in the SNc may counter the MPTP-induced PD symptoms. The gene therapy targeting PKA-RIIβ in the SNc dopaminergic neurons may represent an effective therapeutic way to treat PD.

**Figure 8:**
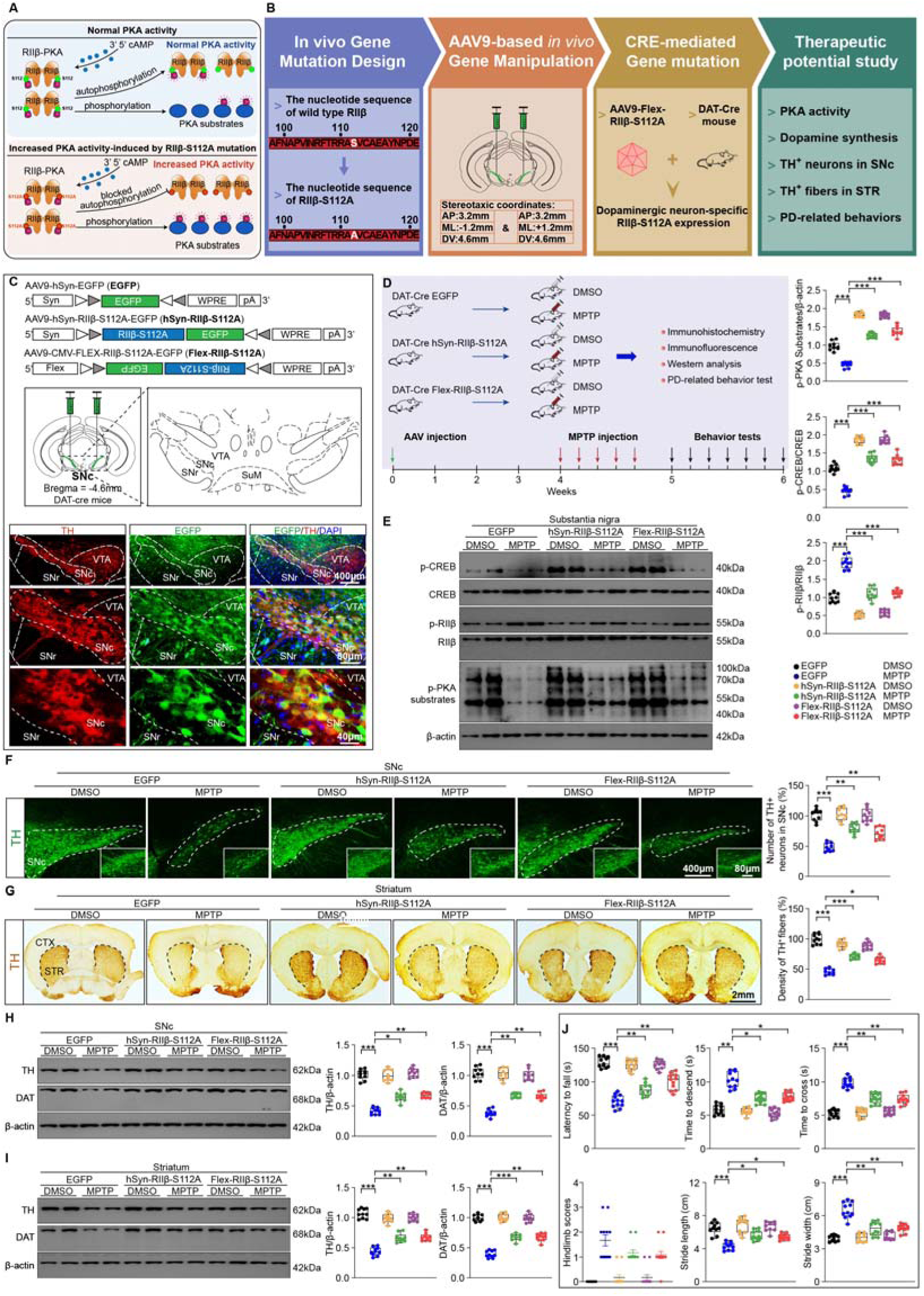
AAV-based gene therapy targeting PKA-RIIβ relieves PD symptoms. **(A)** Schematic diagram of RIIβ-S112A. **(B)** Schematic diagram of the experimental workflow. **(C)** Schematic diagram of AAV and representative fluorescent image showing injection of AAV into the SNc. Scale bars indicate 400 μm, 80 μm and 20 μm, respectively. **(D)** Diagram of the experimental design. **(E)** Representative immunoblots of p-CREB, CREB, p-PKA substrates and β-actin (cropped blot images are shown, see sFigure 10 for full immunoblots) and 1uantification of p-CREB, CREB and p-PKA substrates levels in SNc. Data are mean ± s.e.m.; n = 10 biologically independent animals. Unpaired two tailed Student’s t-tests were used for statistical analyses. \*\*\**p* < 0.001. **(F)** Representative images of TH and unbiased stereological counts of TH-positive neurons in SNc. Data are mean ± s.e.m.; n = 10 biologically independent animals; The two-way ANOVA was used for statistical analysis followed by Tukey’s multiple comparisons test. \*\**p* < 0.01 and \*\*\**p* < 0.001. ns, not significant. Scale bars, 400 μm for low-magnification images and 80 μm for high-magnification images, respectively. **(G)** Representative images of TH staining and density of TH^+^ fibers in striatum. Data are mean ± s.e.m.; n = 12 biologically independent animals; The two-way ANOVA was used for statistical analysis followed by Tukey’s multiple comparisons test. \**p* < 0.05, \*\**p* < 0.01 and \*\*\**p* < 0.001. Scale bar, 2 mm. **(H)** Representative immunoblots of TH, DAT, and β-actin (cropped blot images are shown, see sFigure 10 for full immunoblots) and quantification of TH and DAT levels in SNc. Data are mean ± s.e.m.; n = 9 biologically independent animals. The two-way ANOVA was used for statistical analysis followed by Tukey’s multiple comparisons test. \*\**p* < 0.01 and \*\*\**p* < 0.001. **(I)** Representative immunoblots of TH, DAT, and β-actin (cropped blot images are shown, see sFigure 10 for full immunoblots) and quantification of TH and DAT levels in STR. Data are mean ± s.e.m.; n = 9 biologically independent animals. The two-way ANOVA was used for statistical analysis followed by Tukey’s multiple comparisons test. \*\**p* < 0.01 and \*\*\**p* < 0.001. **(J)** Fall latency from an accelerating rotarod, time to traverse beam apparatus, time to descend pole, Hindlimb clasping reflex score and gait analysis. Data are mean ± s.e.m.; n = 9 biologically independent animals. The two-way ANOVA was used for statistical analysis followed by Tukey’s multiple comparisons test. \**p* < 0.05, \*\**p* < 0.01 and \*\*\**p* < 0.001. ns, not significant.

### An age-related decline of PKA-RIIβ level in SNc dopaminergic neurons underlies PD pathogenesis

As the diagram illustrates in Figure 9, we proposed a simple model describing a dynamic change in PKA activity in the SNc dopaminergic neurons related to PD pathogenesis. In physiological condition, RIIβ subunits preferentially bind to C subunits to form Type-II PKA. The absence or low level of RIIβ subunits results in a compensatory increase of RIα subunits to form Type-I PKA, namely a conversion of Type-II PKA to Type-I PKA. This biochemical adaptation provides an effective mechanism for maintaining the basal PKA activity when RIIβ subunit is absent or lowered. Although there is a compensatory increase of RIα subunits, it was calculated that there is ∼50% loss of R subunits overall as measured by total cAMP-binding capacity. In addition, lower affinity interaction between RIα and C subunits leads to increased degradation of free catalytic subunits. Therefore, PKA activity is reduced in the SNc dopaminergic neurons in PD patients or RIIβ-KO mice. This decreased PKA activity may lead to impaired dopamine synthesis, promoted oxidative stress, and increased apoptosis of dopaminergic neurons, and thus result in spontaneous parkinsonism in old age. PKA-RIIβ subunit in the SNc dopaminergic neurons might be a culprit in PD pathogenesis, and might also be a promising target in gene therapy for PD.

**Figure 9:**
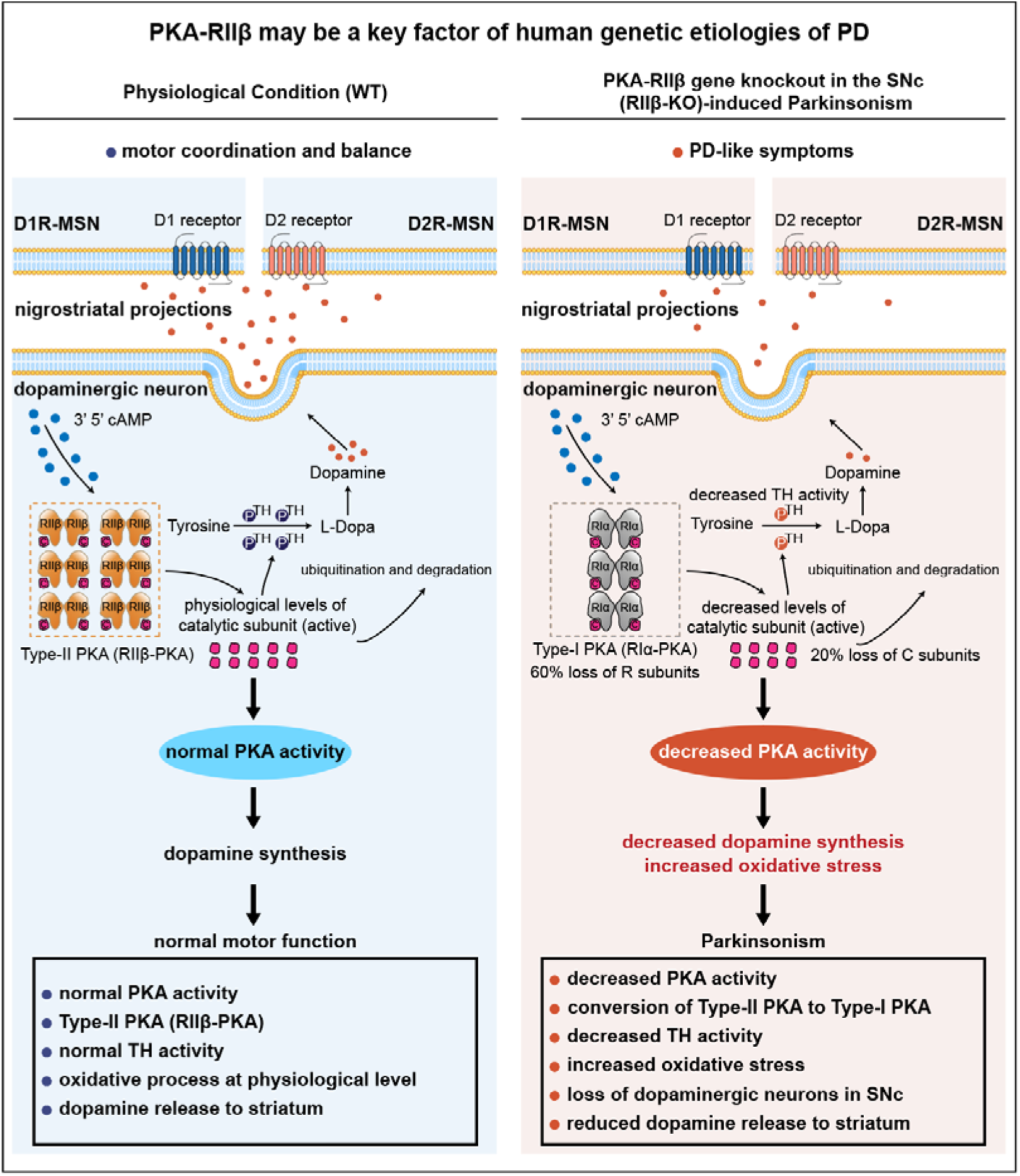
Schematic diagram of an aging-related decline of PKA-RIIβ level in SNc dopaminergic neurons underlies PD pathogenesis.

## Discussion

PKA plays critical roles in the regulation of nigrostriatal dopaminergic function^14–16,20,48,49,64,65^. In the SNc dopaminergic neurons of postmortem specimens from PD patients, there is an inactivation of cAMP response element-binding protein (CREB), a main phosphorylated substrate of PKA^46,66^. In the striatum of demented PD patients, a reduction in cAMP level exists^67^. Pathogenic mutation of PKA regulatory subunit is related to the familial parkinsonism^64^. Aberrant function of PKA substrates is present in PD dopaminergic neuronal cell lines, and this can be rescued by PKA activator^46,47^. PKA activators relive the SNc dopaminergic neuron degeneration and motor symptoms in PD rodent models^18,19^. In this study, by collecting and analyzing the current worldwide human SNc transcriptomic datasets, we discovered that lowered PKA-RIIβ subunit level in the SNc was positively linked to PD pathogenesis. Aged PKA-RIIβ subunit gene knockout mice exhibited remarkable parkinsonism. Taken together, both human and mouse data uncovered that RIIβ-PKA may underlie PD pathogenesis.

PKA regulates tyrosine hydroxylase (TH) for dopamine synthesis. Decreased TH activity is one of the pathological hallmarks of PD^36,37^. PKA phosphorylates TH at Ser 40 to activate TH^39–45^. Decreased PKA activity causes decreased TH activity and reduced dopamine synthesis. Our previous studies revealed that RIIβ-PKA is highly expressed in MSNs of the striatum^49^. Targeted disruption of RIIβ gene expression in the MSNs leads to a decreased PKA activity and aberrant motor function^48,49^. In this study, human SNc transcriptomic bioinformatic analysis revealed a decreased PKA-RIIβ subunit level in the SNc of PD patients. Both our current work and the in situ data of Allen Brain Institute show that RIIβ subunit is expressed in dopaminergic neurons of the SNc. RIIβ deficiency led to decreased PKA activity, lowered TH phosphorylation at Ser 40, reduced dopamine synthesis, and increased oxidative stress in the SNc dopaminergic neurons. Collectively, these findings suggest that decreased PKA-RIIβ subunit level in SNc dopaminergic neurons contributes to PD etiology.

PKA regulates oxidative process of the SNc dopaminergic neurons. The CREB, a key phosphorylated substrate for PKA, can bind the D-loop of mitochondrial genome^68^. Depletion of CREB decreases expression of mitochondria-encoded RNAs of complex I of the electron transport chain^69^. In the familial form of PD, the reduced expression of PINK1 and parkin increases oxidative stress in the SNc dopaminergic neurons^70,71^; transient expression of a constitutively active form of mitochondria-targeted PKA relieves mitochondrial respiratory dysfunction in the PINK1-knockout dopaminergic neurons^17^. PKA phosphorylates subunits of mitochondrial respiratory complex I to facilitate the assembly and enzymatic activity of complex I^72^. PKA inhibits cyclooxygenase (COX) to decrease the ROS production^32,73^. MPTP administration inhibits complex I and increases oxidative stress in the SNc dopaminergic neurons in mice^59^. Rolipram, a PKA activator, relives the degeneration of the SNc dopaminergic neuron in MPTP-induced PD mouse model^19^. In this study, meta-analysis revealed that PKA-RIIβ expression was decreased in the SNc of MPTP-treated mice. In the SNc of MPTP-induced PD mouse model, RIIβ deficiency worsened oxidative stress and aggravated PD-related symptoms. Notably, AAV-mediated gene manipulation targeting PKA-RIIβ subunit for increasing PKA activity effectively mitigated the PD-like symptoms. These findings suggest that RIIβ-PKA may be one of the key regulatory factors that is involved in the oxidative stress of SNc dopaminergic neurons.

Kinase-targeted gene therapy is critical for treating neurodegenerative diseases^74–76^. AAV-mediated dominant-negative c-Jun expression in the striatum protects striatal neurons from neuronal death^77^. In PD mouse models, AAV-delivered N-terminal Myr form of Akt relieves apoptosis of dopaminergic neurons^78,79^. Leucine-rich repeat kinase 2 (LRRK2)-antisense oligonucleotides (ASOs) reduces formation of α-synuclein inclusions^74,80^. In this study, AAV-mediated RIIβ gene mutation for enhancing PKA activity in SNc protected dopaminergic neuronal death and rescued PD behavior in mice. Taken together, these findings suggest that the therapy targeting PKA-RIIβ subunit in SNc dopaminergic neurons may represent an effective therapeutic way to treat PD.

In summary, human SNc transcriptomic analysis reveals that lowered PKA-RIIβ expression level may be associated with PD pathogenesis. In SNc dopaminergic neurons, RIIβ deficiency causes decreased PKA activity, impaired dopamine synthesis, and increased oxidative stress. PKA-RIIβ gene re-expression or AAV-mediated gene manipulation targeting PKA-RIIβ in SNc dopaminergic neurons relieve the parkinsonism of the mice. Thus, these findings indicate that the therapy targeting PKA-RIIβ in SNc dopaminergic neuron may offer a clinically useful way to treat PD, providing a novel insight into PD pathogenesis and therapy.

## Author Contributions

Y.Z. performed the experiment, did the data collection, analyzed the data, made the figures, did the literature search and wrote the paper. B.W., M.Z., D.L., J.L., Y.H. and C.X. participated in the study. R.Z. conceived the study, designed experiments, and wrote and edited the paper. All authors contributed to the study, and reviewed and approved the manuscript for submission. All authors accept full responsibility for the content of this paper.

## Acknowledgments

We thank G. Stanley McKnight (Department of Pharmacology, University of Washington School of Medicine, United States) for kindly providing RIIβ-KO mice and RIIβ ^lox/lox^mice and for helpful discussion. We thank Yuan Zhou (Department of Biomedical Informatics, Center for Noncoding RNA Medicine, School of Basic Medical Sciences, Peking University, China) for bioinformatics support. We thank Bin Zhang (Department of Genetics and Genomic Sciences, Icahn School of Medicine at Mount Sinai, New York, USA) for kind suggestions regarding the MEGENA analysis. We thank Zhuo Huang (Department of Molecular and Cellular Pharmacology, School of Pharmaceutical Sciences, Peking University Health Science Center, Beijing, China) for excellent suggestions regarding the AAV-RIIβ-S112A. We thank Susan S. Taylor (Department of Chemistry and Biochemistry, University of California, San Diego, USA), Ping Zhang (Center for Structural Biology, Center for Cancer Research, National Cancer Institute, Frederick, USA) for helpful suggestions regarding RIIβ holoenzyme structure and molecular dynamics. We thank Yi Rao (Department of Chemical Biology, Peking University, Beijing, China.) for kindly providing D1R-Cre mice. We thank Peng Cao (National Institute of Biological Sciences, Beijing, China.) for kindly providing DAT-cre and A2A-Cre mice. We thank Dennis W. Dickson and Peizhou Jiang (Department of Neuroscience, Mayo Clinic, USA), Elvira De Leonibus (Institute of Genetics and Biophysics, National Research Council, Italy), Shuwen Cao and David G Standaert (Department of Neurology, The University of Alabama at Birmingham, USA), Robert H Edwards (Professor of Neurology and Physiology, UCSF School of Medicine), Marisela Morales (National Institute on Drug Abuse, Neuronal Networks Section, US National Institutes of Health, USA) and Pamela J. McLean (Department of Neuroscience, Mayo Clinic, USA) for their excellent suggestions regarding the AAV delivery. We thank the team of the Annoroad Gene Company (Beijing, China) for support with single-cell sequencing and data analysis, and the team of the Biomedical Sequencing Facility at Novogene for support with next-generation sequencing and data analysis. We thank Hermona Soreq (Department of Biological Chemistry, The Hebrew University of Jerusalem, Jerusalem, Israel), Eva Hedlund (Department of Biochemistry and Biophysics, Stockholm University, Stockholm, Sweden) for discussions about the read length of RNA-seq. We also thank Shigong Zhu (Department of Physiology and Pathophysiology, Peking University, China) for kindly providing access to necessary equipment. We thank Penelope J. Hallett (Neuroregeneration Institute, McLean Hospital/Harvard Medical School, Belmont, USA), Kirk M. Druey (Lung and Vascular Inflammation Section, Laboratory of Allergic Diseases, National Institute of Allergy and Infectious Diseases, National Institutes of Health, Maryland, USA) and Connie Marrase (The Edmond J Safra Program in Parkinson’s disease, Toronto Western Hospital and the University of Toronto, Toronto, Canada) for helpful discussion. We thank Weiguang Zhang, Lihua Qin, Ke Wang (Department of Anatomy, Histology and Embryology, Peking University, China) for technical support.

## References

1 Samii, A., Nutt, J. G. & Ransom, B. R. Parkinson’s disease. Lancet 363, 1783–1793, doi:10.1016/s0140-6736(04)16305-8 (2004).

2 Homayoun, H. Parkinson Disease. Ann Intern Med 169, ITC33–ITC48, doi:10.7326/AITC201809040 (2018).

3 Hopfner, F. et al. beta-adrenoreceptors and the risk of Parkinson’s disease. Lancet Neurol 19, 247–254, doi:10.1016/S1474-4422(19)30400-4 (2020).

4 Obeso, J. A. et al. Missing pieces in the Parkinson’s disease puzzle. Nat Med 16, 653–661, doi:10.1038/nm.2165 (2010).

5 Obeso, J. A., Monje, M. H. G. & Matarazzo, M. Major advances in Parkinson’s disease over the past two decades and future research directions. Lancet Neurol 21, 1076–1079, doi:10.1016/S1474-4422(22)00448-3 (2022).

6 Arkinson, C. & Walden, H. Parkin function in Parkinson’s disease. Science 360, 267–268, doi:10.1126/science.aar6606 (2018).

7 Jankovic, J. & Tan, E. K. Parkinson’s disease: etiopathogenesis and treatment. J Neurol Neurosurg Psychiatry 91, 795–808, doi:10.1136/jnnp-2019-322338 (2020).

8 Gardoni, A. et al. Cerebellar alterations in Parkinson’s disease with postural instability and gait disorders. J Neurol 270, 1735–1744, doi:10.1007/s00415-022-11531-y (2023).

9 Antony, P. M., Diederich, N. J., Kruger, R. & Balling, R. The hallmarks of Parkinson’s disease. FEBS J 280, 5981–5993, doi:10.1111/febs.12335 (2013).

10 Deng, H., Wang, P. & Jankovic, J. The genetics of Parkinson disease. Ageing Res Rev 42, 72–85, doi:10.1016/j.arr.2017.12.007 (2018).

11 Gubellini, P., Picconi, B., Di Filippo, M. & Calabresi, P. Downstream mechanisms triggered by mitochondrial dysfunction in the basal ganglia: from experimental models to neurodegenerative diseases. Biochim Biophys Acta 1802, 151–161, doi:10.1016/j.bbadis.2009.08.001 (2010).

12 Herva, M. E. & Spillantini, M. G. Parkinson’s disease as a member of Prion-like disorders. Virus Res 207, 38–46, doi:10.1016/j.virusres.2014.10.016 (2015).

13 Area-Gomez, E., Guardia-Laguarta, C., Schon, E. A. & Przedborski, S. Mitochondria, OxPhos, and neurodegeneration: cells are not just running out of gas. J Clin Invest 129, 34–45, doi:10.1172/JCI120848 (2019).

14 Kebabian, J. W. & Greengard, P. Dopamine-sensitive adenyl cyclase: possible role in synaptic transmission. Science 174, 1346–1349, doi:10.1126/science.174.4016.1346 (1971).

15 Stoof, J. C. & Kebabian, J. W. Opposing roles for D-1 and D-2 dopamine receptors in efflux of cyclic AMP from rat neostriatum. Nature 294, 366–368, doi:10.1038/294366a0 (1981).

16 Lebel, M., Chagniel, L., Bureau, G. & Cyr, M. Striatal inhibition of PKA prevents levodopa-induced behavioural and molecular changes in the hemiparkinsonian rat. Neurobiol Dis 38, 59–67, doi:10.1016/j.nbd.2009.12.027 (2010).

17 Dagda, R. K. et al. Mitochondrially localized PKA reverses mitochondrial pathology and dysfunction in a cellular model of Parkinson’s disease. Cell Death Differ 18, 1914–1923, doi:10.1038/cdd.2011.74 (2011).

18 Dagda, R. K., Dagda, R. Y., Vazquez-Mayorga, E., Martinez, B. & Gallahue, A. Intranasal Administration of Forskolin and Noopept Reverses Parkinsonian Pathology in PINK1 Knockout Rats. Int J Mol Sci 24, doi:10.3390/ijms24010690 (2022).

19 Yang, L., Calingasan, N. Y., Lorenzo, B. J. & Beal, M. F. Attenuation of MPTP neurotoxicity by rolipram, a specific inhibitor of phosphodiesterase IV. Exp Neurol 211, 311–314, doi:10.1016/j.expneurol.2007.02.010 (2008).

20 Higley, M. J. & Sabatini, B. L. Competitive regulation of synaptic Ca2+ influx by D2 dopamine and A2A adenosine receptors. Nat Neurosci 13, 958–966, doi:10.1038/nn.2592 (2010).

21 Singla, N. & Dhawan, D. K. Influence of zinc on calcium-dependent signal transduction pathways during aluminium-induced neurodegeneration. Mol Neurobiol 50, 613–625, doi:10.1007/s12035-014-8643-7 (2014).

22 Burton, K. A. et al. Type II regulatory subunits are not required for the anchoring-dependent modulation of Ca2+ channel activity by cAMP-dependent protein kinase. Proc Natl Acad Sci U S A 94, 11067–11072, doi:10.1073/pnas.94.20.11067 (1997).

23 Cummings, D. E. et al. Genetically lean mice result from targeted disruption of the RII beta subunit of protein kinase A. Nature 382, 622–626, doi:10.1038/382622a0 (1996).

24 Mellon, P. L., Clegg, C. H., Correll, L. A. & McKnight, G. S. Regulation of transcription by cyclic AMP-dependent protein kinase. Proc Natl Acad Sci U S A 86, 4887–4891, doi:10.1073/pnas.86.13.4887 (1989).

25 Zhang, J. et al. PKA-RIIbeta autophosphorylation modulates PKA activity and seizure phenotypes in mice. Commun Biol 4, 263, doi:10.1038/s42003-021-01748-4 (2021).

26 Liu, Y., Chen, J., Fontes, S. K., Bautista, E. N. & Cheng, Z. Physiological and pathological roles of protein kinase A in the heart. Cardiovasc Res 118, 386–398, doi:10.1093/cvr/cvab008 (2022).

27 Cheung, J. et al. Structural insights into mis-regulation of protein kinase A in human tumors. Proc Natl Acad Sci U S A 112, 1374–1379, doi:10.1073/pnas.1424206112 (2015).

28 Cadd, G. & McKnight, G. S. Distinct patterns of cAMP-dependent protein kinase gene expression in mouse brain. Neuron 3, 71–79, doi:10.1016/0896-6273(89)90116-5 (1989).

29 Niswender, C. M. et al. Cre recombinase-dependent expression of a constitutively active mutant allele of the catalytic subunit of protein kinase A. Genesis 43, 109–119, doi:10.1002/gene.20159 (2005).

30 Amieux, P. S. et al. Compensatory regulation of RIalpha protein levels in protein kinase A mutant mice. J Biol Chem 272, 3993–3998, doi:10.1074/jbc.272.7.3993 (1997).

31 Willis, B. S., Niswender, C. M., Su, T., Amieux, P. S. & McKnight, G. S. Cell-type specific expression of a dominant negative PKA mutation in mice. PLoS One 6, e18772, doi:10.1371/journal.pone.0018772 (2011).

32 Bouchez, C. & Devin, A. Mitochondrial Biogenesis and Mitochondrial Reactive Oxygen Species (ROS): A Complex Relationship Regulated by the cAMP/PKA Signaling Pathway. Cells 8, doi:10.3390/cells8040287 (2019).

33 Bogacka, I., Ukropcova, B., McNeil, M., Gimble, J. M. & Smith, S. R. Structural and functional consequences of mitochondrial biogenesis in human adipocytes in vitro. J Clin Endocrinol Metab 90, 6650–6656, doi:10.1210/jc.2005-1024 (2005).

34 Di Benedetto, G., Gerbino, A. & Lefkimmiatis, K. Shaping mitochondrial dynamics: The role of cAMP signalling. Biochem Biophys Res Commun 500, 65–74, doi:10.1016/j.bbrc.2017.05.041 (2018).

35 Kostic, M. et al. PKA Phosphorylation of NCLX Reverses Mitochondrial Calcium Overload and Depolarization, Promoting Survival of PINK1-Deficient Dopaminergic Neurons. Cell Rep 13, 376–386, doi:10.1016/j.celrep.2015.08.079 (2015).

36 Nagatsu, T. Catecholamines and Parkinson’s disease: tyrosine hydroxylase (TH) over tetrahydrobiopterin (BH4) and GTP cyclohydrolase I (GCH1) to cytokines, neuromelanin, and gene therapy: a historical overview. J Neural Transm (Vienna) 131, 617–630, doi:10.1007/s00702-023-02673-y (2024).

37 Smolders, S. & Van Broeckhoven, C. Genetic perspective on the synergistic connection between vesicular transport, lysosomal and mitochondrial pathways associated with Parkinson’s disease pathogenesis. Acta Neuropathol Commun 8, 63, doi:10.1186/s40478-020-00935-4 (2020).

38 Gopinath, A. et al. DAT and TH expression marks human Parkinson’s disease in peripheral immune cells. NPJ Parkinsons Dis 8, 72, doi:10.1038/s41531-022-00333-8 (2022).

39 Dunkley, P. R. & Dickson, P. W. Tyrosine hydroxylase phosphorylation in vivo. J Neurochem 149, 706–728, doi:10.1111/jnc.14675 (2019).

40 Sura, G. R., Daubner, S. C. & Fitzpatrick, P. F. Effects of phosphorylation by protein kinase A on binding of catecholamines to the human tyrosine hydroxylase isoforms. J Neurochem 90, 970–978, doi:10.1111/j.1471-4159.2004.02566.x (2004).

41 Daubner, S. C., Le, T. & Wang, S. Tyrosine hydroxylase and regulation of dopamine synthesis. Arch Biochem Biophys 508, 1–12, doi:10.1016/j.abb.2010.12.017 (2011).

42 Roskoski, R., Jr. & Ritchie, P. Phosphorylation of rat tyrosine hydroxylase and its model peptides in vitro by cyclic AMP-dependent protein kinase. J Neurochem 56, 1019–1023, doi:10.1111/j.1471-4159.1991.tb02023.x (1991).

43 Fujisawa, H. & Okuno, S. Regulatory mechanism of tyrosine hydroxylase activity. Biochem Biophys Res Commun 338, 271–276, doi:10.1016/j.bbrc.2005.07.183 (2005).

44 Dunkley, P. R., Bobrovskaya, L., Graham, M. E., von Nagy-Felsobuki, E. I. & Dickson, P. W. Tyrosine hydroxylase phosphorylation: regulation and consequences. J Neurochem 91, 1025–1043, doi:10.1111/j.1471-4159.2004.02797.x (2004).

45 Stoop, J., Douma, E. H., van der Vlag, M., Smidt, M. P. & van der Heide, L. P. Tyrosine hydroxylase phosphorylation is under the control of serine 40. J Neurochem 167, 376–393, doi:10.1111/jnc.15963 (2023).

46 Chalovich, E. M., Zhu, J. H., Caltagarone, J., Bowser, R. & Chu, C. T. Functional repression of cAMP response element in 6-hydroxydopamine-treated neuronal cells. J Biol Chem 281, 17870–17881, doi:10.1074/jbc.M602632200 (2006).

47 Dagda, R. K. & Das Banerjee, T. Role of protein kinase A in regulating mitochondrial function and neuronal development: implications to neurodegenerative diseases. Rev Neurosci 26, 359–370, doi:10.1515/revneuro-2014-0085 (2015).

48 Brandon, E. P. et al. Defective motor behavior and neural gene expression in RIIbeta-protein kinase A mutant mice. J Neurosci 18, 3639–3649, doi:10.1523/JNEUROSCI.18-10-03639.1998 (1998).

49 Zheng, R. et al. Deficiency of the RIIbeta subunit of PKA affects locomotor activity and energy homeostasis in distinct neuronal populations. Proc Natl Acad Sci U S A 110, E1631–1640, doi:10.1073/pnas.1219542110 (2013).

50 Yang, L. & McKnight, G. S. Hypothalamic PKA regulates leptin sensitivity and adiposity. Nat Commun 6, 8237, doi:10.1038/ncomms9237 (2015).

51 Gong, S. et al. Targeting Cre recombinase to specific neuron populations with bacterial artificial chromosome constructs. J Neurosci 27, 9817–9823, doi:10.1523/JNEUROSCI.2707-07.2007 (2007).

52 Gong, S. et al. A gene expression atlas of the central nervous system based on bacterial artificial chromosomes. Nature 425, 917–925, doi:10.1038/nature02033 (2003).

53 Zhuang, X., Masson, J., Gingrich, J. A., Rayport, S. & Hen, R. Targeted gene expression in dopamine and serotonin neurons of the mouse brain. J Neurosci Methods 143, 27–32, doi:10.1016/j.jneumeth.2004.09.020 (2005).

54 Zhao, M. et al. The DJ1-Nrf2-STING axis mediates the neuroprotective effects of Withaferin A in Parkinson’s disease. Cell Death Differ 28, 2517–2535, doi:10.1038/s41418-021-00767-2 (2021).

55 Jackson-Lewis, V. & Przedborski, S. Protocol for the MPTP mouse model of Parkinson’s disease. Nat Protoc 2, 141–151, doi:10.1038/nprot.2006.342 (2007).

56 Song, W. M. & Zhang, B. Multiscale Embedded Gene Co-expression Network Analysis. PLoS Comput Biol 11, e1004574, doi:10.1371/journal.pcbi.1004574 (2015).

57 Rempe, D. et al. Synapsin I Cre transgene expression in male mice produces germline recombination in progeny. Genesis 44, 44–49, doi:10.1002/gene.20183 (2006).

58 Wang, B. et al. RIIbeta-PKA in GABAergic Neurons of Dorsal Median Hypothalamus Governs White Adipose Browning. Adv Sci (Weinh) 10, e2205173, doi:10.1002/advs.202205173 (2023).

59 Dionisio, P. A., Amaral, J. D. & Rodrigues, C. M. P. Oxidative stress and regulated cell death in Parkinson’s disease. Ageing Res Rev 67, 101263, doi:10.1016/j.arr.2021.101263 (2021).

60 Zhang, P. et al. Single Turnover Autophosphorylation Cycle of the PKA RIIbeta Holoenzyme. PLoS Biol 13, e1002192, doi:10.1371/journal.pbio.1002192 (2015).

61 Roskoski, R., Jr. A historical overview of protein kinases and their targeted small molecule inhibitors. Pharmacol Res 100, 1–23, doi:10.1016/j.phrs.2015.07.010 (2015).

62 Zhang, P. et al. Structure and allostery of the PKA RIIbeta tetrameric holoenzyme. Science 335, 712–716, doi:10.1126/science.1213979 (2012).

63 Isensee, J. et al. PKA-RII subunit phosphorylation precedes activation by cAMP and regulates activity termination. J Cell Biol 217, 2167–2184, doi:10.1083/jcb.201708053 (2018).

64 Wong, T. H. et al. PRKAR1B mutation associated with a new neurodegenerative disorder with unique pathology. Brain 137, 1361–1373, doi:10.1093/brain/awu067 (2014).

65 Parisiadou, L. et al. LRRK2 regulates synaptogenesis and dopamine receptor activation through modulation of PKA activity. Nat Neurosci 17, 367–376, doi:10.1038/nn.3636 (2014).

66 Xu, X. et al. CREB Inactivation by HDAC1/PP1gamma Contributes to Dopaminergic Neurodegeneration in Parkinson’s Disease. J Neurosci 42, 4594–4604, doi:10.1523/JNEUROSCI.1419-21.2022 (2022).

67 Nishino, N., Kitamura, N., Hashimoto, T. & Tanaka, C. Transmembrane signalling systems in the brain of patients with Parkinson’s disease. Rev Neurosci 4, 213–222, doi:10.1515/revneuro.1993.4.2.213 (1993).

68 Ryu, H., Lee, J., Impey, S., Ratan, R. R. & Ferrante, R. J. Antioxidants modulate mitochondrial PKA and increase CREB binding to D-loop DNA of the mitochondrial genome in neurons. Proc Natl Acad Sci U S A 102, 13915–13920, doi:10.1073/pnas.0502878102 (2005).

69 Lee, J. et al. Mitochondrial cyclic AMP response element-binding protein (CREB) mediates mitochondrial gene expression and neuronal survival. J Biol Chem 280, 40398–40401, doi:10.1074/jbc.C500140200 (2005).

70 Funayama, M. et al. Familial Parkinsonism with digenic parkin and PINK1 mutations. Mov Disord 23, 1461–1465, doi:10.1002/mds.22143 (2008).

71 Ishihara-Paul, L. et al. PINK1 mutations and parkinsonism. Neurology 71, 896–902, doi:10.1212/01.wnl.0000323812.40708.1f (2008).

72 Signorile, A. et al. cAMP/PKA Signaling Modulates Mitochondrial Supercomplex Organization. Int J Mol Sci 23, doi:10.3390/ijms23179655 (2022).

73 Helling, S. et al. Phosphorylation and kinetics of mammalian cytochrome c oxidase. Mol Cell Proteomics 7, 1714–1724, doi:10.1074/mcp.M800137-MCP200 (2008).

74 Sun, J. & Roy, S. Gene-based therapies for neurodegenerative diseases. Nat Neurosci 24, 297–311, doi:10.1038/s41593-020-00778-1 (2021).

75 Perez, D. I., Gil, C. & Martinez, A. Protein kinases CK1 and CK2 as new targets for neurodegenerative diseases. Med Res Rev 31, 924–954, doi:10.1002/med.20207 (2011).

76 Kawahata, I. & Fukunaga, K. Protein Kinases and Neurodegenerative Diseases. Int J Mol Sci 24, doi:10.3390/ijms24065574 (2023).

77 Hayley, S. et al. Regulation of dopaminergic loss by Fas in a 1-methyl-4-phenyl-1,2,3,6-tetrahydropyridine model of Parkinson’s disease. J Neurosci 24, 2045–2053, doi:10.1523/JNEUROSCI.4564-03.2004 (2004).

78 Burke, R. E. Inhibition of mitogen-activated protein kinase and stimulation of Akt kinase signaling pathways: Two approaches with therapeutic potential in the treatment of neurodegenerative disease. Pharmacol Ther 114, 261–277, doi:10.1016/j.pharmthera.2007.02.002 (2007).

79 Ries, V. et al. Oncoprotein Akt/PKB induces trophic effects in murine models of Parkinson’s disease. Proc Natl Acad Sci U S A 103, 18757–18762, doi:10.1073/pnas.0606401103 (2006).

80 Zhao, H. T. et al. LRRK2 Antisense Oligonucleotides Ameliorate alpha-Synuclein Inclusion Formation in a Parkinson’s Disease Mouse Model. Mol Ther Nucleic Acids 8, 508–519, doi:10.1016/j.omtn.2017.08.002 (2017).

