## Supplemental figures and tables for "Age-related decline of PKA-RIIβ level in SNc dopaminergic neurons underlies PD pathogenesis"

**Supplemental figure 1**

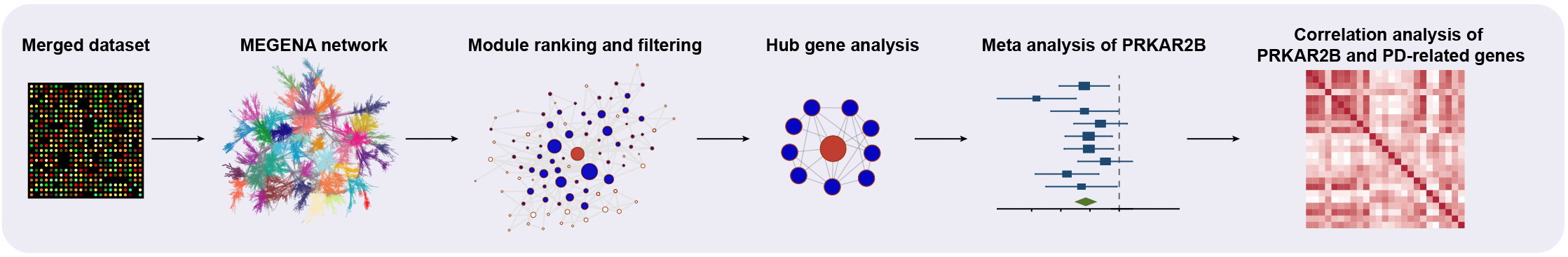

**Figure S1: Schematic representation of the bioinformatics method used in this study. MEGENA, multiscale embedded gene co-expression network analysis**

**Supplemental figure 2**

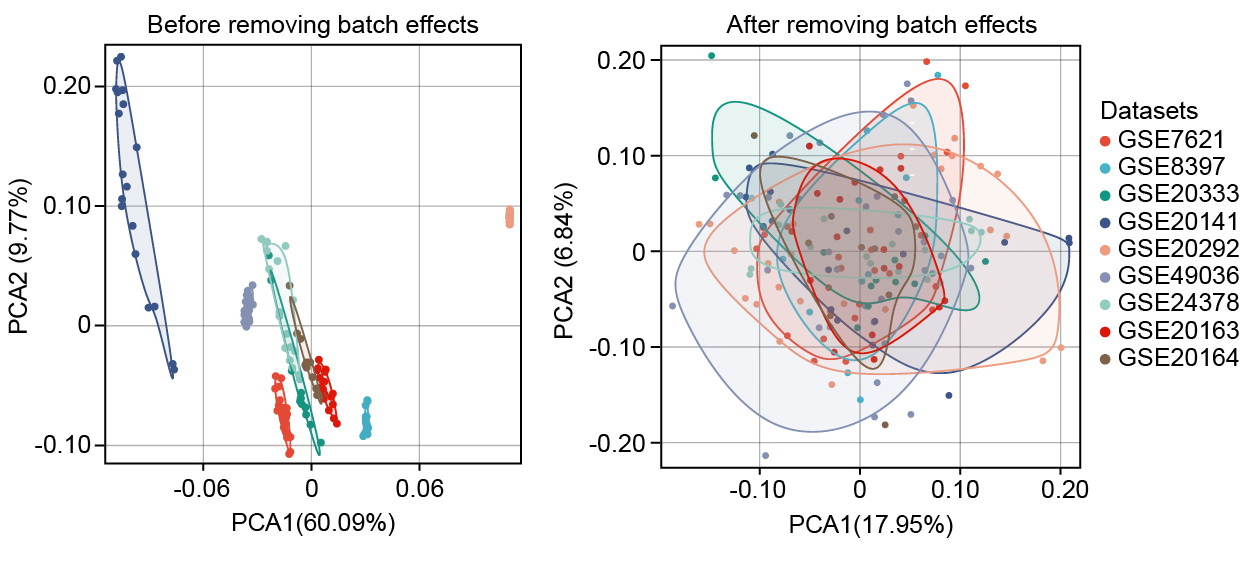

**Figure S2: Principle component analysis (PCA) plot before and after removing batch effects using SVA comBat for 9 datasets**

**Supplemental figure 3**

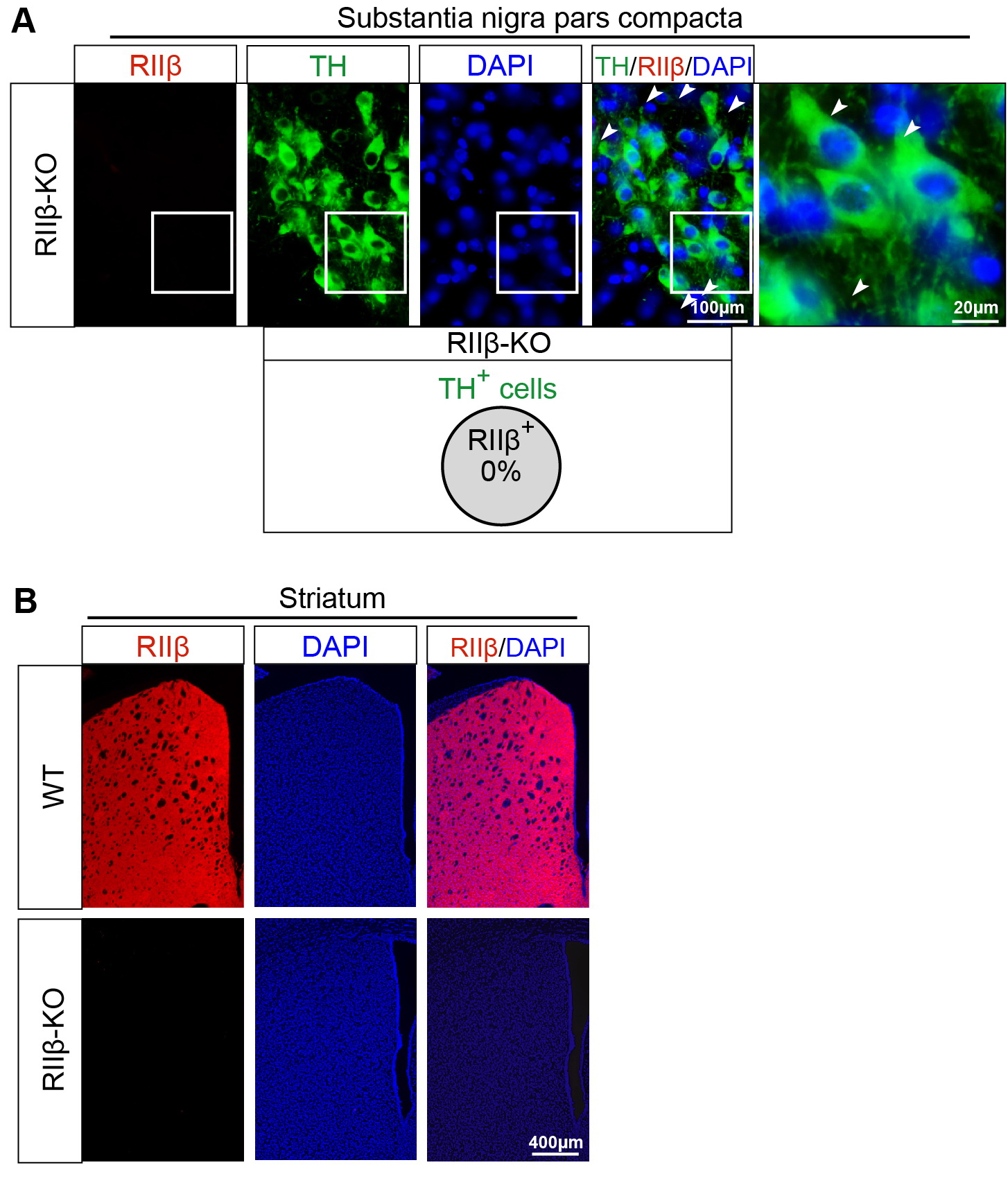

**Figure S3: RIIβ protein was observed in SNc dopaminergic neurons and RIIβ protein was knockout in RIIβ^-/-^ mice**

(A) Representative double immunostaining of RIIβ with TH in SNc of RIIβ^-/-^ mice. Percentage of double-positive for TH (green) and RIIβ (red) was shown in the bottom. Scale bar, 100 μm for low-magnification images and 20 μm for high-magnification images, respectively.

(B) Representative immunostaining of RIIβ in STR of WT and RIIβ^-/-^ mice. Scale bar, 400 μm.

**Supplemental figure 4**

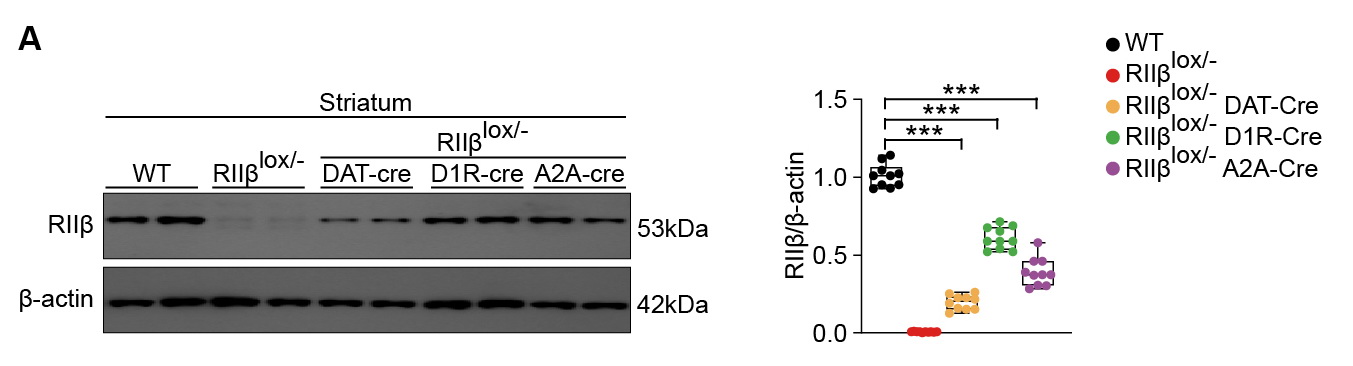

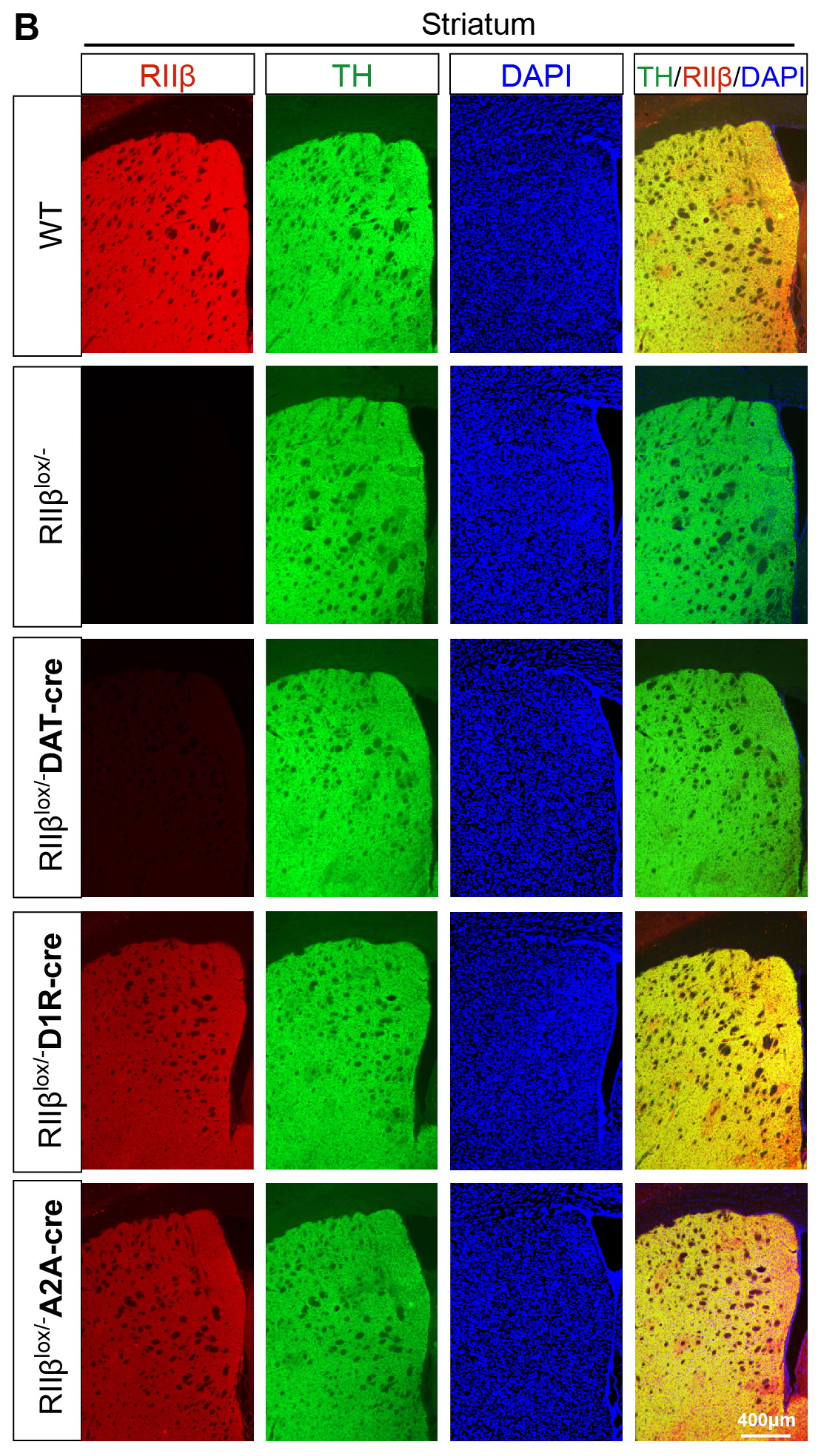

**Figure S4: RIIβ reexpression in SNc dopaminergic neurons, D1R striatal MSNs or D2R striatal MSNs**

(A) Representative immunoblots of RIIβ and β-actin (cropped blot images are shown, see sFigure 9 for full immunoblots) and quantification of RIIβ levels in the STR. Data are mean ± s.e.m.; n = 9 biologically independent animals. The two-way ANOVA was used for statistical analysis followed by Tukey’s multiple comparisons test. ****p* < 0.001.

(B) Representative immunostaining of RIIβ in STR of WT, RIIβ^lox/-^, RIIβ^lox/-^ DAT-Cre, RIIβ^lox/-^ D1R-Cre and RIIβ^lox/-^ A2A-Cre mice. Scale bar, 400 μm.

**Supplemental figure 5**

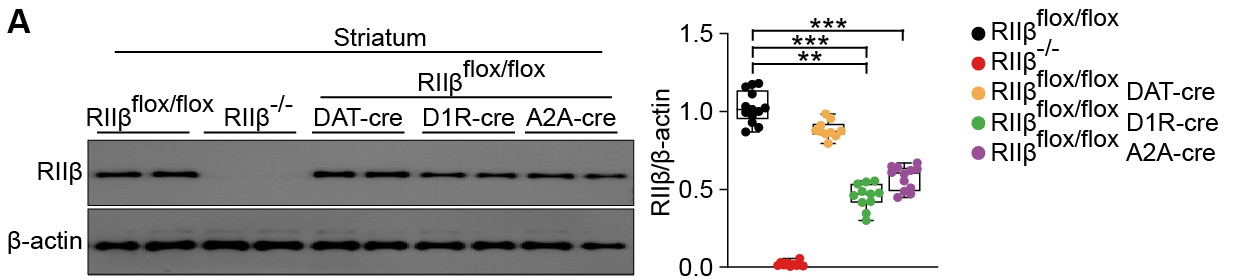

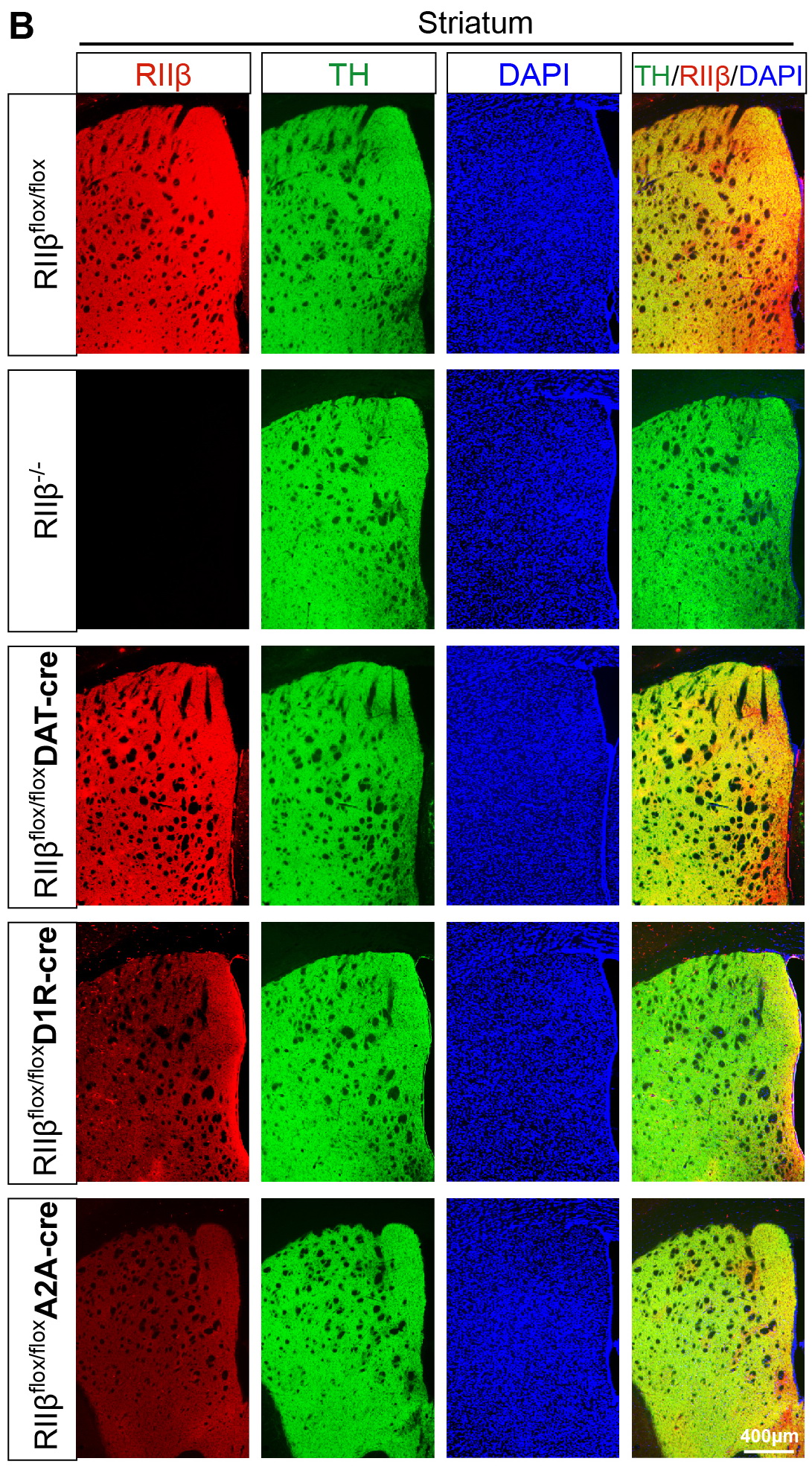

**Figure S5: RIIβ was specifically knocked out in SNc dopaminergic neurons, D1R striatal MSNs or D2R striatal MSNs**

(A) Representative immunoblots of RIIβ and β-actin (cropped blot images are shown, see sFigure 9 for full immunoblots) and quantification of RIIβ levels in STR. Data are mean ± s.e.m.; n = 9 biologically independent animals. The two-way ANOVA was used for statistical analysis followed by Tukey’s multiple comparisons test. ***p* < 0.01 and ****p* < 0.001.

(B) Representative immunostaining of RIIβ in STR of RIIβ^flox/flox^, RIIβ^-/-^, RIIβ^flox/flox^ DAT-cre, RIIβ^flox/flox^ D1R-cre and RIIβ^flox/flox^ A2A-cre mice. Scale bar, 400 μm.

**Supplemental figure 6**

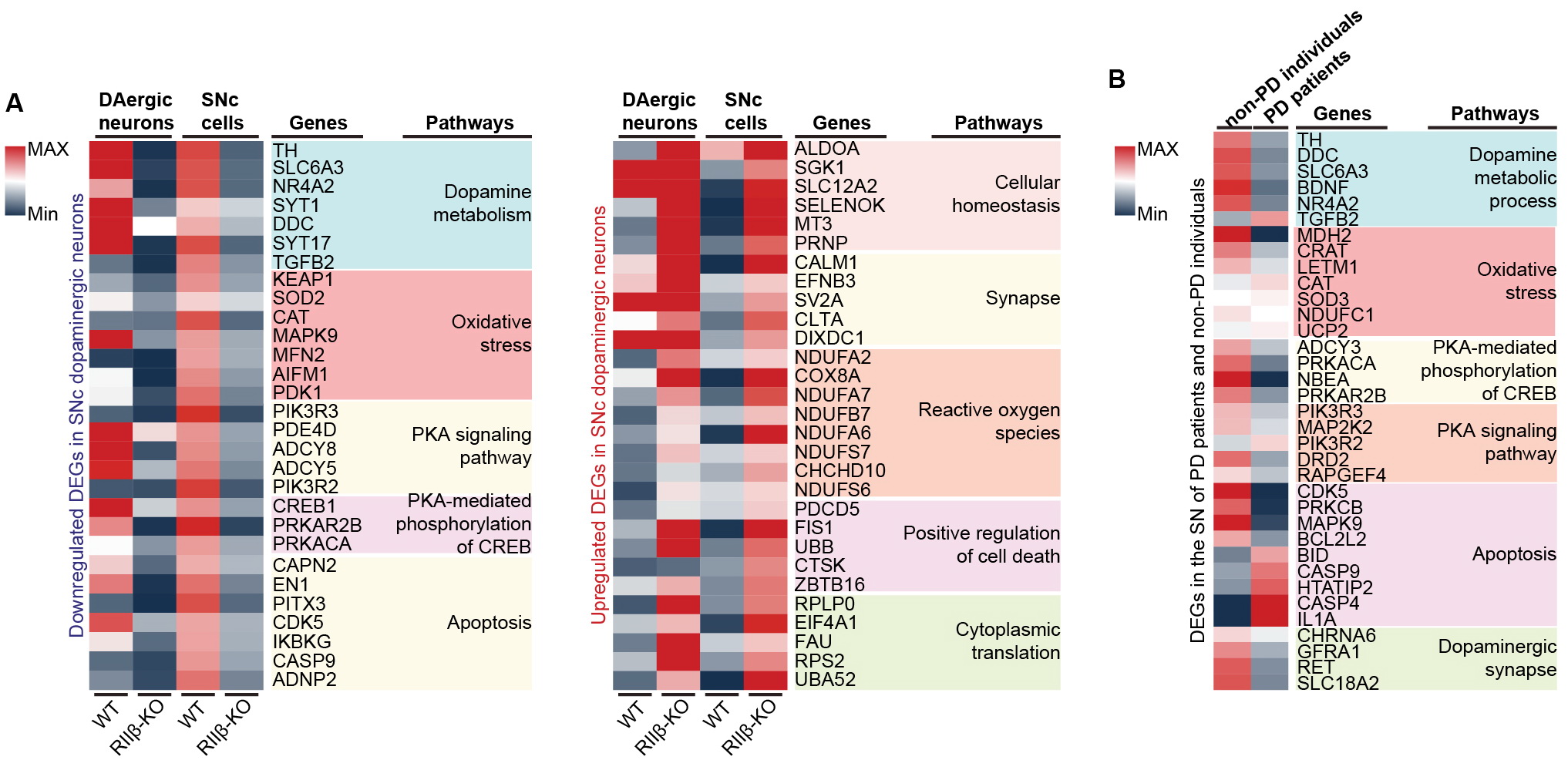

**Figure S6: Decreased PKA activity and dopamine synthesis in SNc dopaminergic neurons of RIIβ-KO mice**

(A) Heatmap of DEGs at both single-nucleus RNA sequencing (dopaminergic neurons) resolution in the SNc of human (WT and RIIβ-KO mouse). (*p* < 0.05 with unpaired two-tailed Student’s t tests). (B) Heatmap of DEGs in the SN of human (PD patients and non-PD individuals). (*p* < 0.05 with unpaired two-tailed Student’s t tests).

**Supplemental figure 7**

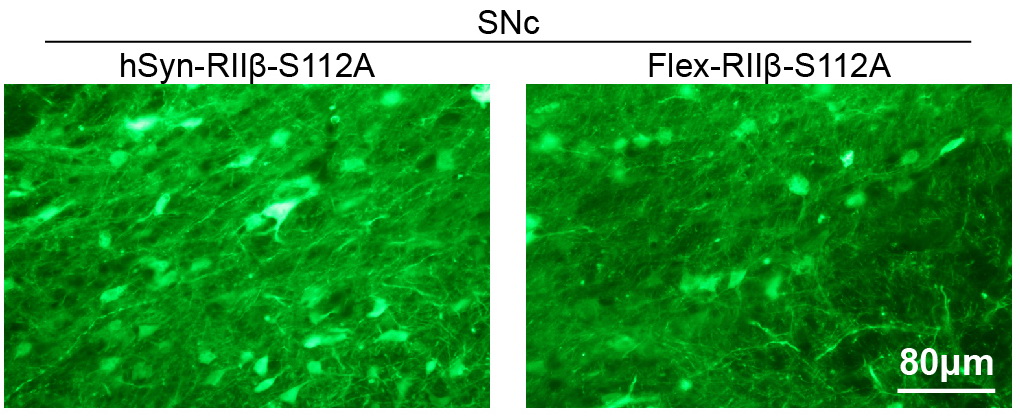

**Figure S7: Mice transfected with AAV were transcardially perfused on day 21 after AAV injection. Scale bar, 80 μm, green indicating AAV-transduced dopaminergic neurons.**

**Supplemental figure 8**

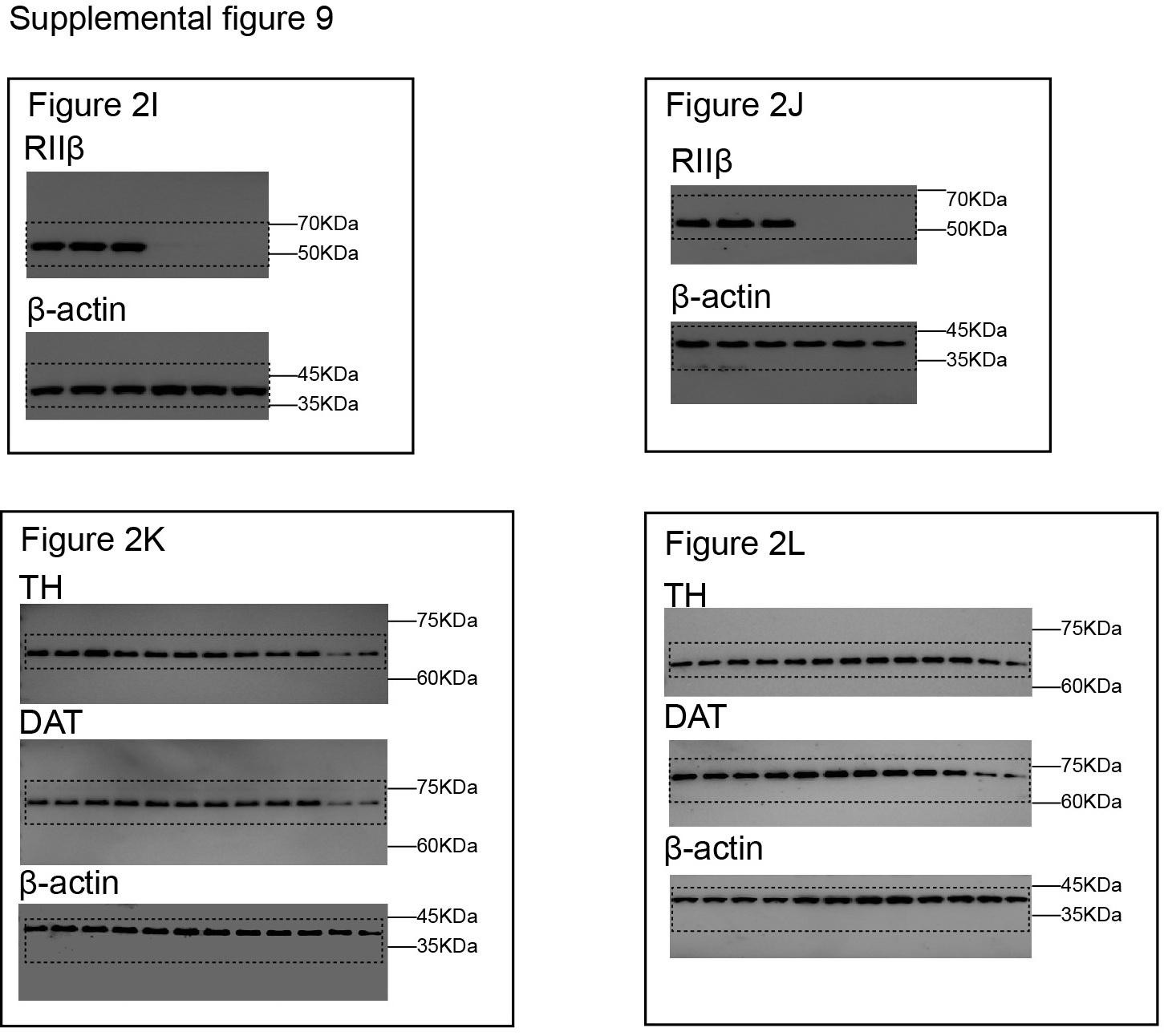

**Figure S8: Original full western blot images of Figure 2.**

**Supplemental figure 9**

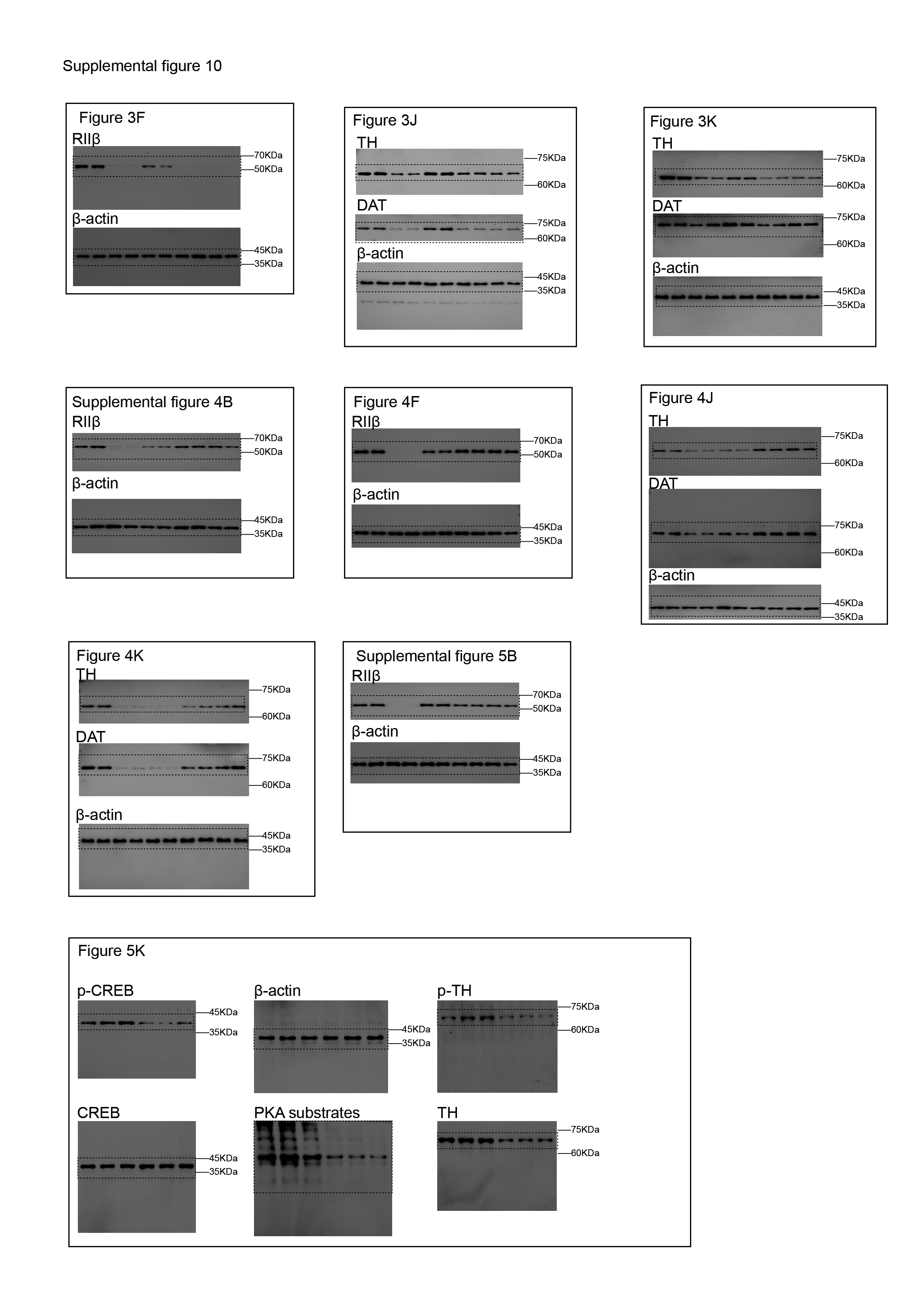

**Figure S9: Original full western blot images of Figure 3, Figure 4, Figure S4, Figure S5 and Figure 5.**

**Supplemental figure 10**

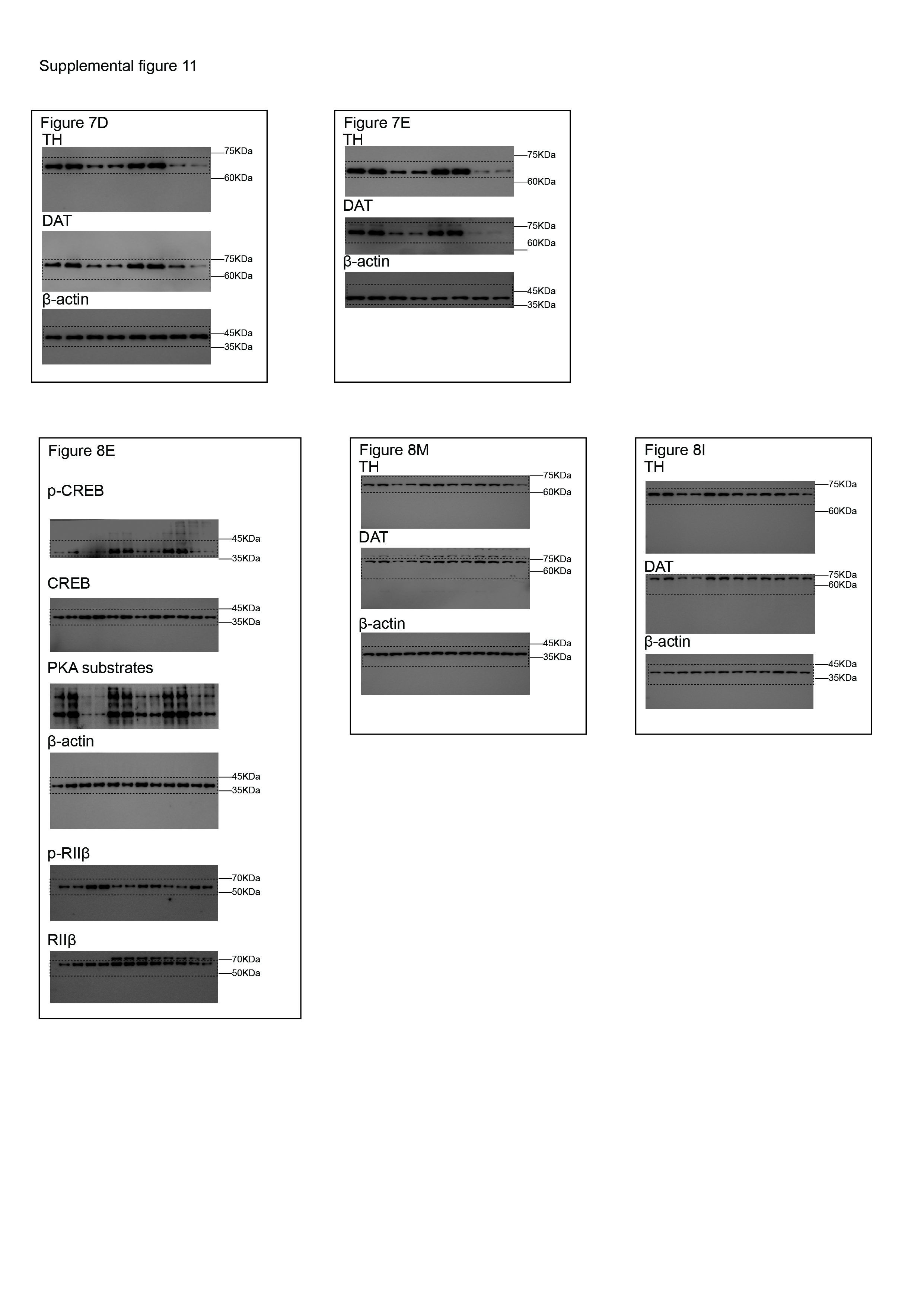

**Figure S10: Original full western blot images of Figure 7 and Figure 8.**

**Supplementary Table 1: Antibodies used in this study.**

| **Antibodies** | **Source/Cat. No.** | **Host** | **Dilution** |
| --- | --- | --- | --- |
| Tyrosine Hydroxylase (TH) | Millipore (AB152) | Rabbit | 1:1,000 (WB)  1:400 (IHC, IF) |
| p-TH (Ser40) | Cell Signaling (2791s) | Rabbit | 1:1,000 (WB) |
| Dopamine transporter (DAT) | Bioss (bs-1714R) | Rabbit | 1:1,000 (WB) |
| β-actin | sigma (A5316) | Mouse | 1:5000 (WB) |
| PKA-RIIβ | BD Biosciences (610625) | Mouse | 1:2000 (WB)  1:400 (IF) |
| PKA-RIIβ (pS114) | BD Biosciences (612550) | Mouse | 1:1000 (WB) |
| CREB | Cell Signaling (9197s) | Rabbit | 1:1,000 (WB) |
| p-CREB (Ser133) | Cell Signaling (9198s) | Rabbit | 1:1,000 (WB) |
| p-PKA Substrate (RRXS*/T*) | Cell Signaling (9624) | Rabbit | 1:1,000 (WB) |

| **Supplementary Table 2. Human postmortem brain studies of PD in public domain** | | | | |  |
| --- | --- | --- | --- | --- | --- |
| **GEO ID** | **Brain Region** | **non-PD** | **PD** | **Platform** | **Pre-Processed** |
| GSE7621 | substantia nigra | 9 | 16 | Affymetrix Human Genome U133 Plus 2.0 Array | MAS5 |
| GSE8397 | substantia nigra | 7 | 9 | Affymetrix Human Genome U133A Array | RMA |
| GSE20333 | substantia nigra | 6 | 6 | Affymetrix Human HG-Focus Target Array | MAS5 |
| GSE20141 | substantia nigra | 8 | 10 | Affymetrix Human Genome U133 Plus 2.0 Array | MAS5 |
| GSE20292 | substantia nigra | 15 | 11 | Affymetrix Human Genome U133A Array | MAS5 |
| GSE49036 | substantia nigra | 8 | 20 | Affymetrix Human Genome U133 Plus 2.0 Array | GC-RMA |
| GSE24378 | substantia nigra | 9 | 8 | Affymetrix Human X3P Array | MAS5 |
| GSE20163 | substantia nigra | 8 | 9 | Affymetrix Human Genome U133A Array | MAS5 |
| GSE20164 | substantia nigra | 5 | 6 | Affymetrix Human Genome U133A Array | MAS5 |
|  | total | 75 | 95 |  |  |
